# Constitutive activity of ionotropic glutamate receptors via hydrophobic substitutions in the ligand-binding domain

**DOI:** 10.1101/2023.08.03.551817

**Authors:** Sandra Seljeset, Oksana Sintsova, Yuhong Wang, Hassan Y. Harb, Timothy Lynagh

**Affiliations:** Michael Sars Centre, University of Bergen, 5008 Bergen, Norway; Concept Life Sciences Limited, Frith Knoll Road, Chapel-en-le-Frith, High Peak, SK23 0PG, United Kingdom

**Keywords:** binding site, glutamate, iGluR, ion channels, LBD, ligand-gated ion channels, neurotransmitter receptors, NMDA receptor, Placozoa, spontaneous activity

## Abstract

Neurotransmitter ligands electrically excite neurons by activating ionotropic glutamate receptor (iGluR) ion channels. Knowledge of the iGluR amino acid residues that dominate ligand-induced activation would enable the prediction of function from sequence. We therefore explored the molecular determinants of activity in rat NMDA-type iGluRs (NMDA receptors), complex heteromeric iGluRs comprising two glycine-binding GluN1 and two glutamate-binding GluN2 subunits, using amino acid sequence analysis, mutagenesis, and electrophysiology. We find that a broadly conserved aspartate residue controls both ligand potency and channel activity, to the extent that certain substitutions at this position bypass the need for ligand-binding in GluN1 subunits, generating NMDA receptors activated solely by glutamate. Furthermore, we identify a homomeric iGluR from the placozoan *Trichoplax adhaerens* that has utilized native mutations of this crucial residue to evolve into a leak channel that is inhibited by neurotransmitter binding, pointing to a dominant role of this residue throughout the iGluR superfamily.

## INTRODUCTION

Neurons rapidly signal to each other via chemical synapses, where neurotransmitter released by one cell diffuses across a narrow synaptic cleft and binds to ligand-gated ion channels (LGICs), or “ionotropic receptors”, on the postsynaptic cell membrane ^1,2^. LGICs are membrane proteins with extracellular ligand-binding domains (LBDs) and a membrane-spanning channel domain. Ligand-binding in the LBD triggers conformational changes that open (i.e. “gate”) the channel within milliseconds, mediating rapid ion flux across the cell membrane. This depolarizes the postsynaptic neuron in the case of excitatory, Na^+^-permeable channels, typified by ionotropic glutamate receptors (iGluRs), the major excitatory receptor in the mammalian brain ^3,4^.

iGluRs are tetramers, formed by the assembly of four homologous subunits around a central channel pore, with each subunit contributing its own extracellular LBD ^4^. The iGluR superfamily is broad, but most members bind the transmitters glutamate or glycine, and the superfamily can be divided into several families based on functional differences and phylogenetic relationships ^5^. Mammalian synapses typically contain iGluRs from two families: α-amino-3-hydroxy-5-methyl-4-isoxazolepropionate-sensitive iGluRs (AMPA receptors); and N-methyl-D-aspartate-sensitive iGluRs (NMDA receptors). Like primitive iGluRs from bacteria and early-branching animals ^6,7^, AMPA receptor subunits GluA1-GluA4 readily express heterologously as homo-tetramers that rapidly activate and deactivate in response to glutamate, and *in vivo*, AMPA receptors conduct mostly Na^+^and K^+^ ^8^. In contrast, NMDA receptors are obligate hetero-tetramers, comprising two glycine- and D-serine-binding GluN1 subunits and two glutamate-binding GluN2 subunits, thus requiring two different ligands for channel opening ^9^. GluN1/GluN2 NMDA receptors are important “coincidence detectors” as they carry relatively long-lasting Na^+^ and Ca^2+^ currents *in vivo* only after channel block by Mg^2+^ ions is relieved by membrane depolarization ^10^. Glycine- and D-serine-binding GluN3 subunits also assemble with GluN1 into GluN1/GluN3 NMDA receptors, which occur more sporadically in mammalian brain and respond to glycine with small, rapidly desensitizing currents^11,12^.

Channel gating requires ligand binding in the clamshell-shaped LBD, during which the lower lobe of the clamshell moves upward, away from the cell membrane. This opens the channel by pulling on the channel-lining α-helices via short peptide segments that link the LBD to the channel ^13–15^. LBD clamshell closure is a broadly conserved feature of glutamate-gated homo-tetramers, glycine-gated homo-tetramers, and glutamate and glycine-binding hetero-tetramers alike, as inferred by X-ray crystallographic structures of diverse iGluRs ^6,15,16^. However, several aspects of LBD activity deserve investigation. From a biophysical perspective where we seek to understand channel gating mechanisms, the contributions of four distinct LBDs to the gating of a shared, central channel remains unclear ^17–19^, and structural studies suggest that clamshell closure may not drive channel gating the same way in different iGluR families ^20,21^. And from a biological and pharmacological perspective, predicting iGluR function from amino acid sequence remains difficult: despite high-resolution structural data, we don’t entirely understand which LBD residues dominate ligand recognition and channel activity ^22^.

We therefore extensively investigated the role of one particular amino acid residue in iGluR activation. We focused on it because of its broad conservation as either a glutamate or aspartate residue in diverse iGluRs, its position in the middle of the LBD clamshell, and its previous implication in both ligand recognition and LBD conformational change. We began our study utilizing the glycine-binding GluN1 subunit of hetero-tetrameric NMDA receptors, where the residue in question is an aspartate, D732. X-ray crystallography studies have shown that the GluN1-D732 side chain orients away from the ligand-binding site in inactive receptor states but toward glycine in active receptor states ^15,23^, and a mutagenesis study showed that even the relatively subtle D732E substitution drastically reduces the potency of glycine in activating GluN1 subunits ^24^. We explored the role of this residue in NMDA receptor activity and also in a distantly related homomeric iGluR, using mutagenesis, electrophysiology, and amino acid sequence analysis of diverse iGluRs. This uncovered a remarkably dominant contribution of this position to both ligand recognition and channel activity, not only in heteromeric NMDA receptors but also in distantly related homomeric iGluRs.

## RESULTS

### GluN1 D732L and -F substitutions yield NMDA receptors activated solely by glutamate

We first investigated the contribution of D732 to glycine-induced activation of GluN1 subunits by replacing it with amino acid residues of varying physico-chemical properties (Fig. 1A). We then measured currents through GluN1-mutant/GluN2A-wildtype receptors heterologously expressed in *Xenopus laevis* oocytes, in response to increasing glycine concentrations in the presence of 100 µM glutamate. Compared to wildtype (WT) channels, glycine potency was decreased 1,000-10,000-fold in channels with small side chains (D732A, D732S), with a longer carboxylate side chain (D732E), or with an isosteric but amide side chain (D732N) at the GluN1-D732 position (Fig. 1B), consistent with previous work on GluN1/GluN2B receptors ^24^. We also observed large decreases in glycine potency with β-branched hydrophobic D732V and D732I substitutions (Fig. 1A,B). Longer hydrophobic substitutions, however, had very different effects. At GluN1-D732L- and GluN1-D732F-containing receptors, we saw no glycine concentration-dependence of ligand-gated currents, with large currents already activated by 100 nM glycine/100 µM glutamate (Fig. 1B). Similarly, large currents were activated in mutant GluN1-D732L/GluN2A-WT receptors when 100 nM through 100 μM D-Serine was applied in the presence of 100 µM glutamate (green in Fig. 1B). Thus, the additional lower LBD lobe interactions possible with D-serine relative to glycine ^15^ make no apparent difference to GluN1-D732L/GluN2A-WT activity.

**Figure 1.**
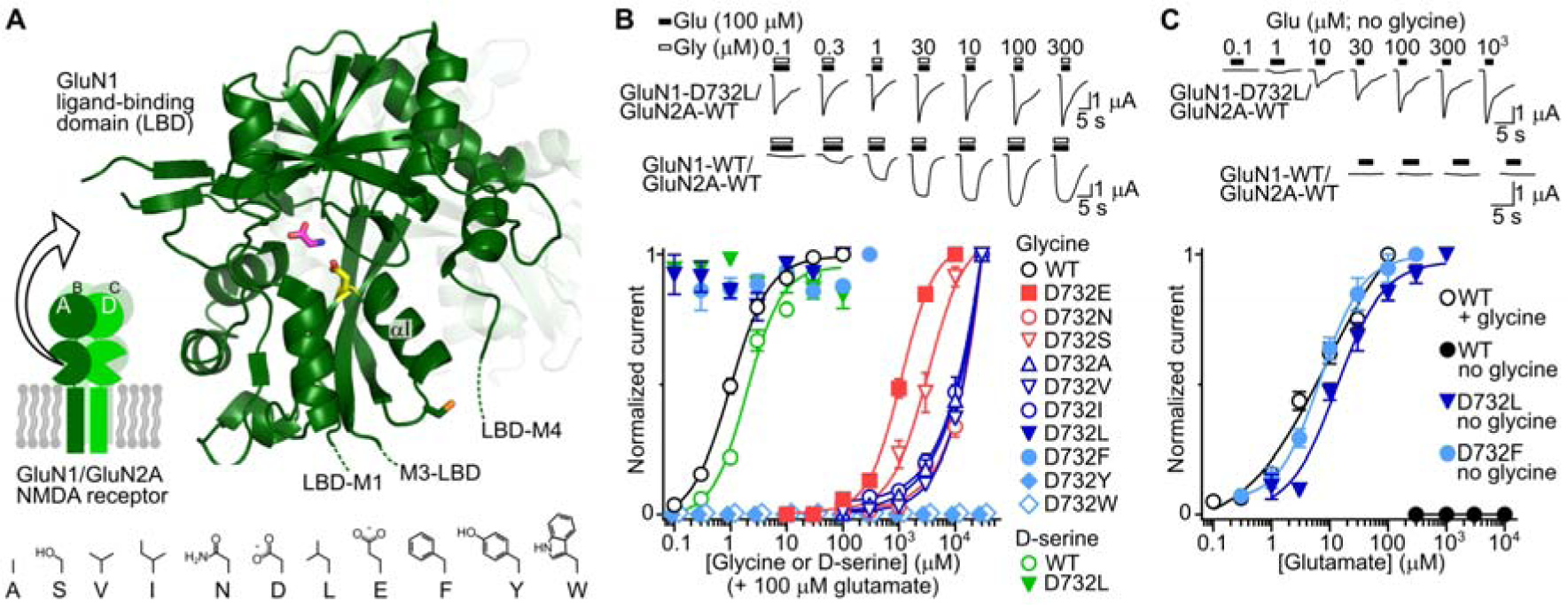
Certain hydrophobic side chains at the GluN1 732 position yield NMDA receptors activated solely by glutamate. (**A**) Mid-left, cartoon hetero-tetrameric NMDA receptor showing GluN1 subunits (dark green), GluN2A subunits (light green), and surrounding cell membrane (grey). Note that GluN1 subunits occupy positions “A” and “C” in the tetramer when counted clockwise from above ^25^. Main, Bound glycine (magenta) and GluN1 D732 side chain (yellow) in previously published X-ray structure of NMDA receptor ligand-binding domain (PDB 4NF8). αI helix labelled and following C744 side chain shown. Dashed lines indicate approximate position of LBD-channel domain linkers. Bottom, amino acid side chain 2D structures. (**B**) Upper, example glycine+glutamate-gated currents in Xenopus laevis oocytes expressing indicated GluN1/GluN2A receptors. Lower, glycine concentration-current response relationships (data points: mean ± SEM, n = 6-7) for GluN1/GluN2A receptors carrying indicated GluN1 subunits. (**C**) Upper, example glutamate-gated currents in the absence of glycine in oocytes expressing indicated GluN1/GluN2A receptors. Lower, corresponding glutamate concentration-current response relationships (data points: mean ± SEM, n = 6-7).

Reasoning that this effect may derive from solely glutamate-gated currents in GluN1-D732L- and GluN1-D732F-containing receptors, we tested responses of these receptors to increasing concentrations of glutamate in the absence of co-agonists. Indeed, glutamate alone elicited concentration-dependent currents in GluN1-D732L/GluN2A-WT and GluN1-D732F/GluN2A-WT receptors (Fig. 1C), with concentration-response relationships (glutamate EC_50_ 12.8 ± 2.0 µM and 7.1 ± 1.4 µM, respectively, both n = 6) that were very similar to that of glutamate at WT receptors in the presence of 100 µM glycine (glutamate EC_50_ 4.8 ± 0.5, n = 11, Fig. 1C). Tyrosine and tryptophan substitution in the same position did not induce the same effect, as we observed no glutamate-or glycine and glutamate-gated currents at oocytes injected with GluN1-D732Y or GluN1-D732W together with GluN2A-WT RNAs (Fig. 1B). These data show that most substitutions at the GluN1-732 position decrease glycine potency, but leucine and phenylalanine substitutions alter GluN1 activity in such a way that leads to single-mutant NMDA receptors activated solely by glutamate.

### D732L and -F mutations render GluN1 subunits constitutively active but LBD conformation is unclear

Solely glutamate-gated currents at GluN1-D732L and D732F-containing receptors could be due to the L and F side chains conferring glutamate sensitivity on GluN1 LBDs, such that glutamate now binds to GluN1 and GluN2A subunits to activate the mutant receptors. Alternatively, the L and F side chains might induce an active GluN1 LBD conformation in the absence of ligand, such that glutamate activates the mutant receptors by only binding to GluN2A subunits, reflected in previous studies that locked GluN1 LBDs in an active state via introduced disulfide bonds ^26,27^. To distinguish between these possibilities, we measured glutamate-gated currents in receptors comprising GluN1 double-mutant R523K/D732L and GluN2A-WT subunits. R523 is a highly conserved arginine residue in the LBD of diverse iGluRs, and in GluN1 and GluN2A subunits its guanidino side chain binds the α-carboxylate of glycine and glutamate, respectively, and its mutation to lysine greatly decreases agonist potency (Fig. 2A and ref.^28^). Receptors containing GluN1-R523K/D732L were potently activated by glutamate in the absence of glycine much like those containing GluN1-D732L (Fig. 2A, GluN1-R523K/D732L EC_50_ 3.5 ± 1.7 µM, n = 3), whereas receptors containing single-mutant GluN1-R523K were not activated by glycine and glutamate below 3 mM (n = 5, Fig. 2A). Assuming that glutamate does not bind to GluN1-R523K LBDs in an unforeseen binding mode, this suggests that glutamate activates GluN1-D732L-containing receptors by binding strictly to GluN2A-WT subunits, and D732L and D732F substitutions render GluN1 subunits constitutively active in a manner similar to glycine-binding.

**Figure 2.**
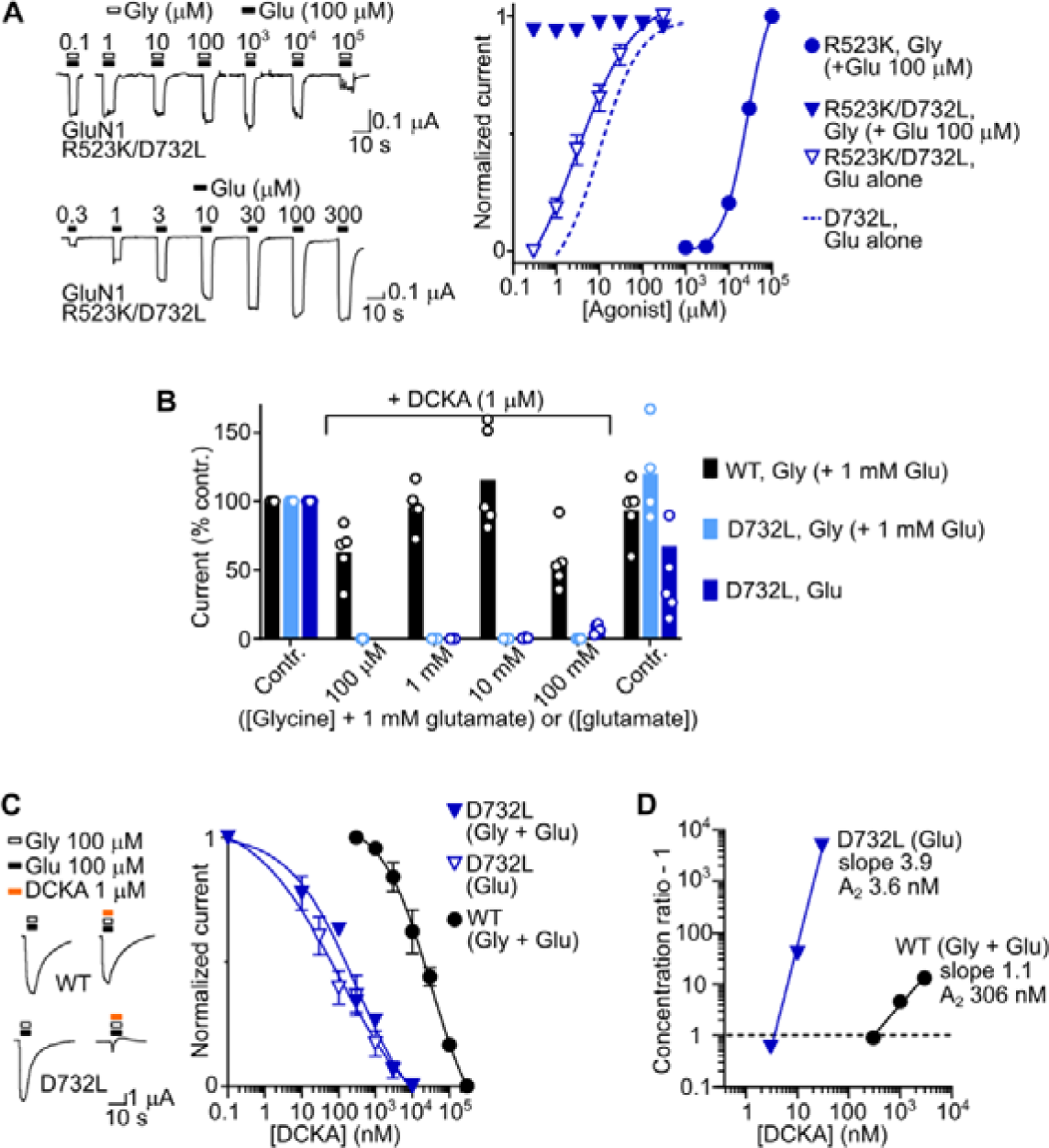
GluN1 D732L subunits do not require ligand binding for activation. (**A**) Left, example ligand-gated currents in Xenopus laevis oocytes expressing NMDA receptors comprising GluN1-R523K/D732L and GluN2A-WT subunits. Right, concentration-current response relationships (mean ± SEM, n = 3-5) for receptors carrying indicated GluN1 subunits together with WT GluN2A subunits. (**B**) Mean (columns) and individual data points (n = 4-6) of glycine (+ 1 mM glutamate)-gated currents in GluN1-WT and GluN1-D732L-containing receptors and glutamate-gated currents in GluN1-D732L-containing receptors in the absence (control, “contr.”) or presence of 1 μM 5,7-dichlorokynurenic acid (DCKA). (**C**) Left, example and right, mean (± SEM, n = 4) current in the presence of increasing DCKA concentrations in oocytes expressing indicated GluN1 subunits and WT GluN2A subunits. (**D**) Schild analysis for indicated GluN1 mutants. Full analysis, Fig. S2.

Seeking complementary evidence that glutamate does not bind to mutant GluN1 subunits, we measured competition for the GluN1 LBD between glutamate and 5,7-dichlorokynurenic acid (DCKA), a GluN1-selective competitive antagonist ^29^. Neither glutamate at concentrations up to 100 mM, nor glycine up to 100 mM in the presence of 1 mM glutamate were able to relieve DCKA-induced inhibition of GluN1-D732L/GluN2A-WT receptors, in contrast to WT receptors, where even 100 μM glycine relieved more than 50% DCKA-induced inhibition (Fig. 2B). This shows that DCKA binding is intact in GluN1-D732 mutants and is further indicative that activation of GluN1-D732L/GluN2A WT-receptors by glutamate is not via glutamate binding to mutant GluN1 subunits.

As mutant GluN1-D732L subunits are constitutively active, iGluR activation typically involves LBD clamshell closure, and DCKA binding stabilizes an LBD clamshell-open conformation ^15^, it is perhaps surprising that DCKA effectively inhibits GluN1-D732L-containing receptors. We explored this further by measuring DCKA potency at mutant receptors (Fig. 2C). DCKA potency was in fact drastically increased in mutant receptors (IC_50_ 440 ± 120 nM in the presence of glycine and glutamate, n = 4; IC_50_ 280 ± 230 nM in the presence of glutamate alone, n = 4) relative to WT receptors (IC_50_ 24.9 ± 5.2 µM, n = 4). Schild analysis ^30^ confirmed that whereas DCKA and glycine compete for the GluN1 site in WT receptors (slope ≈ 1, Fig. 2D and Fig. S2), DCKA inhibition of mutant receptors is not classically competitive (slope ≈ 4, Fig. 2D and Fig. S2). Whether increased DCKA potency in GluN1-D732L subunits derives from the now non-competitive nature of the inhibition, or from the introduction of a favourable hydrophobic interaction with the dichlorobenzene moiety of the inhibitor, is unclear. But the high DCKA potency would suggest that the constitutively active GluN1-D732L subunit is, unexpectedly, not due to a permanently clamshell-closed LBD in the mutant. This may reflect the fact that extent of LBD closure is poorly correlated with agonist efficacy in GluN1 subunits, in contrast to AMPA receptor GluA2 subunits ^21^.

In order to understand whether the glycine-insensitive GluN1-D732L subunit is in a constantly activated state or occupies a different conformation that may reflect an alternative to typical channel gating, we compared the kinetics of WT receptor and GluN1-D732L-containing receptor inhibition by the open-channel blocker MK-801, which can be used to evaluate maximum open probability of NMDARs ^26,31^. We observed very similar kinetics of inhibition of WT and mutant receptors (Fig. 3A), indicating similar open probability in solely glutamate-gated GluN1-D732L-containing receptors and glutamate and glycine-gated WT receptors. This reflects unchanged maximum open probability in solely glutamate-gated NMDARs with disulfide-locked GluN1 LBDs assayed by single channel recordings ^27^. This suggests that the GluN1-D732L subunit is in a constantly activated state.

**Figure 3.**
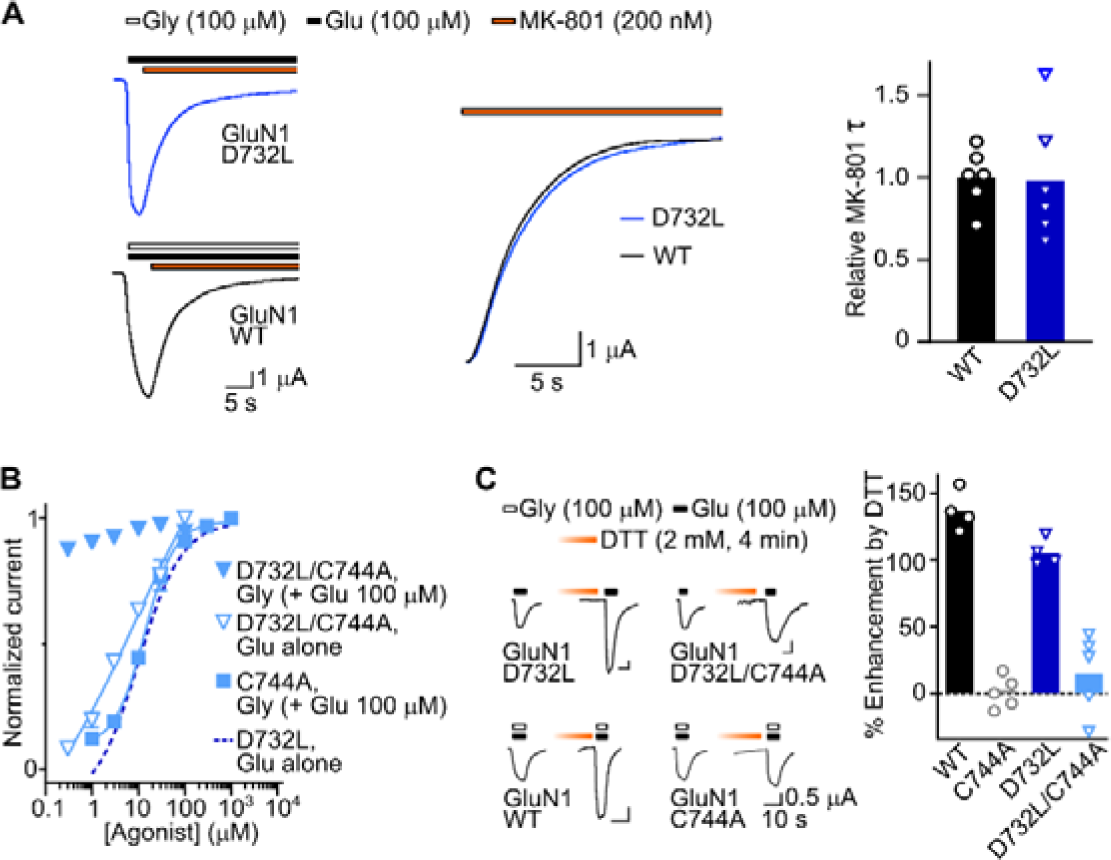
Effect of disulfide removal on GluN1-D732L activity. (**A**) Left, Example recordings of inhibition of ligand-gated currents by MK-801 in oocytes expressing NMDA receptors comprising indicated mutant GluN1 subunits and WT GluN2A subunits. Mid, Magnified view. Right, individual (points, n = 6) and mean (columns) MK-801-induced current decay “τ”, normalized to WT mean τ. (**B**) Concentration-current response relationships (data points mean ± SEM, n = 3-5) for receptors carrying indicated GluN1 subunits together with WT GluN2A subunits. (**C**) Left, example ligand-gated currents, before and after dithiothreitol (DTT) treatment. Right, mean (columns) and individual data points (n = 4-5) % enhancement of current amplitude by DTT.

As another means of ruling out alternative channel gating in GluN1-D732L receptors, we tested if redox modulation was intact. Although most energy of channel activation derives from the LBD pulling on the upper M3 helix, the gating machinery also includes the peptide segment linking the C-terminal end of the LBD to membrane-spanning helix 4 (LBD-M4 linker) ^13^. LBD and LBD-M4 linker conformation is confined by a C744—C798 disulfide, just four helical turns after D732, and disruption of this disulfide enhances channel gating ^32^. If channels of GluN1-D732L-containing receptors are activated much like WT receptors, we expected that current enhancement via reduction of the C744—C798 via dithiothreitol (DTT) would reflect that of WT receptors, and that conversely, removal of the disulfide via mutagenesis would leave the constitutive activity of mutant GluN1 subunits intact. Indeed, GluN1-D732L/C744A-containing receptors were activated by glutamate alone (EC_50_ = 21 ± 4 μM, n = 4, Fig. 3B). Moreover, DTT-induced enhancement of glutamate-gated currents was intact in single-mutant GluN1-D732L-containing receptors, and this enhancement was abolished by the addition of the C744A mutation (Fig. 3C). Thus, we conclude that the GluN1-D732L subunit is constantly in a typically activated channel state, despite the fact the LBD retains sensitivity to DCKA.

### D732-αI interactions are replaced by Q536-αI interactions during GluN1 activation by glycine

Our results with mutant receptors suggest that the 732 side chain controls ligand potency and channel activity, and we therefore asked how D732 mediates these roles in WT channels. D732 movements during GluN1 activation can be inferred from previously published X-ray structures of the isolated GluN1 LBD in different states: competitive antagonist-bound inactive; ligand-free inactive; and glycine-bound active (Fig. 4A). The D732 side chain is positioned to accept hydrogen (H) bonds from A734 and V735 of the αI helix in inactive states, but swings centrally in the glycine-bound state to interact with the agonist amine (Fig. 4A). We reasoned that replacing A734 and V735 NH groups with an H bond acceptor would remove D732—αI interactions specific to inactive states and bias D732 toward its active position, resulting in either increased glycine potency or constitutive GluN1 activity similar to GluN1-D732L and D732F. We therefore measured activity of NMDA receptors in which GluN1 A734 and V735 were replaced by their α-hydroxy acid analogues, lactic acid (Aah) and 2-hydroxy-3-methylbutyric acid (Vah) via the nonsense suppression method (Fig. S1). However, neither the A734Aah nor the V735Vah substitution caused constitutive activity or an increase in glycine potency (Fig. 4B), indicating that the removal of single D732—αI interactions is not sufficient to bias the GluN1 subunit toward the active state.

**Figure 4.**
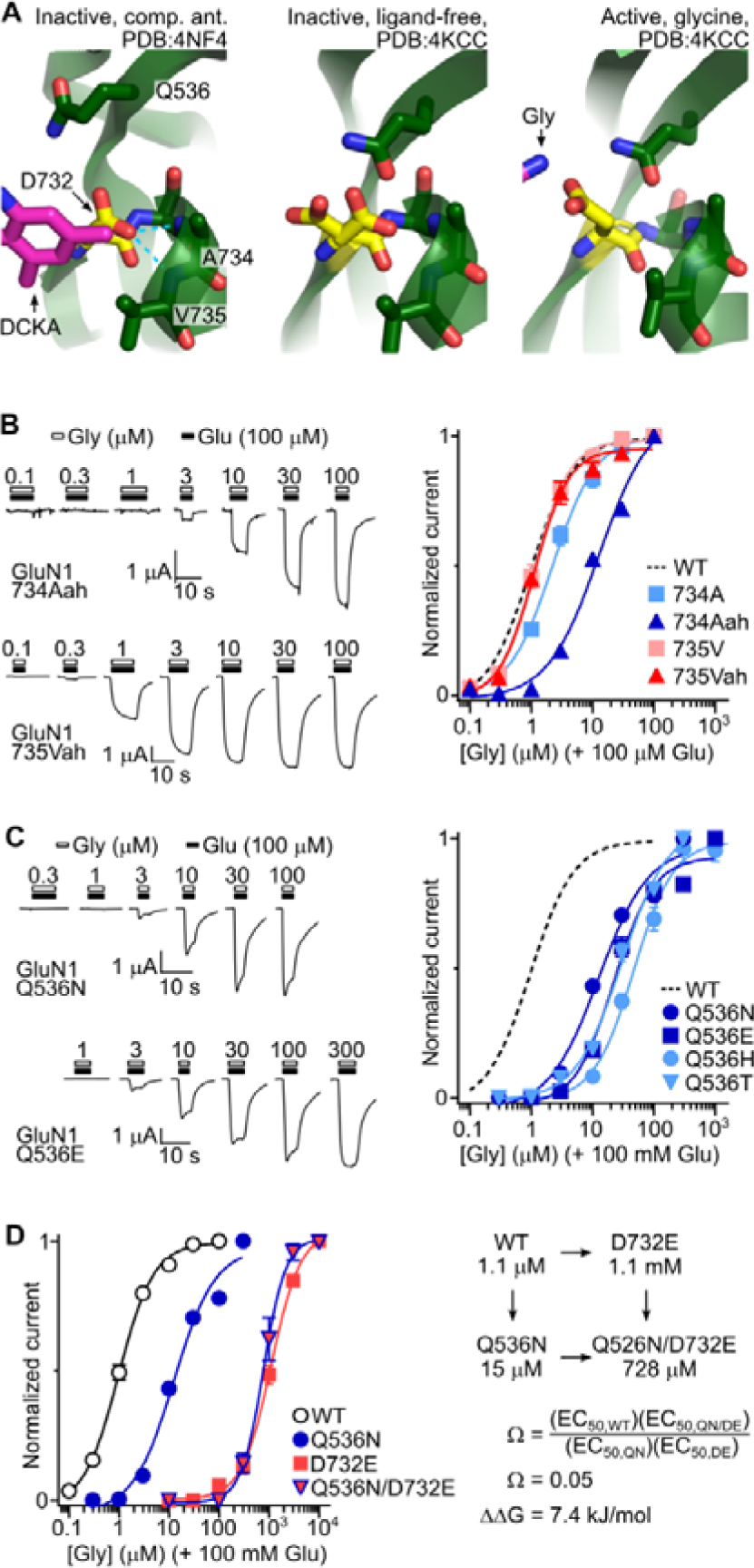
Q536 replaces D732 in H bond accepting from A734 during LBD activation. (**A**) D732 in indicated GluN1 X-ray structures. Comp. ant., competitive antagonist, 5,7-dichlorokynurenic acid (DCKA). Dashed lines, putative H-bonds. (**B,C**) Left, example ligand-gated currents in Xenopus laevis oocytes expressing NMDA receptors comprising indicated mutant GluN1 subunits and WT GluN2A subunits. Aah, lactic acid. Vah, 2-hydroxy-3-methylbutyric acid. Right, concentration-current response relationships (data points mean ± SEM, n = 5-7) for receptors carrying indicated GluN1 subunits together with WT GluN2A subunits. 734A and 735V in (**B**) refer to controls in which “WT” residues were incorporated via nonsense suppression (Fig. S1). (**D**) Left, mean ± SEM (n = 4) normalized glycine-gated current at GluN1-Q536N/D732E-containing receptors (others repeated from above), and right, mutant cycle analysis yielding coupling parameter, Ω, and interaction energy ΔΔG =-RTln(Ω).

The A734Aah substitution in fact caused a fourfold *decrease* in glycine potency (EC_50_ values 10.6 ± 0.6 µM, n = 6 for A734Aah and 2.4 ± 0.3 µM, n = 6, for WT in the nonsense suppression method; Fig. 4B, Fig. S1). Although this may be due to local conformational changes due to altered main chain structure, we considered that this decrease in potency may result from the loss of some interaction with the A734 NH that is conducive to the active state. We therefore re-examined the glycine-bound structure and observed that the GluN1 Q536 side chain from β-strand 12 in the LBD hinge region essentially moves 6 Å from the competitive antagonist-bound state and 0.4 Å from the ligand-free state toward both the A734 NH and the D732 COOH (Fig. 4A). Thus, the decreased glycine potency in GluN1-A734Aah may be via the loss of an A734 amine—Q536 amide bond that stabilizes the active LBD. We tested this interpretation by measuring the effects of conventional Q536N, -E, -T, and -H substitutions, which decrease the length of the amide side chain (Q536N), remove a hydrogen bond donor (Q536E), or replace the amide side chain with smaller or larger polar moieties (Q536T, Q536H). We observed 15-fold (Q536N, EC_50_ = 15 ± 0.6 μM, n = 5) to 50-fold (Q536H, EC_50_ = 51 ± 5 μM, n = 5) decreases in glycine potency in these mutants (Fig. 4C), indicating that the amide moiety and to a slightly smaller extent the length of the Q536 side chain are significant determinants of glycine-induced activation.

Combined with existing X-ray structures, this suggests that D732 is important for glycine-induced activation of the WT LBD in part because glycine binding to one carboxylate oxygen of D732 exposes the αI-A734 amine to the Q536 amide oxygen. This conformation is in turn stabilized by the other D732 carboxylate oxygen interacting with the Q536 amide nitrogen, as implied by previous X-ray structural studies ^15,23^. If an interaction between Q536 and D732 were energetically important for receptor activation, the effects of their mutations should be non-additive ^33^. We therefore tested glycine potency at double-mutant GluN1-Q536N/D732E-containing receptors and observed non-additive changes in EC_50_, with a strong coupling value, Ω, of 0.05 (Fig. 4D). This deviation of Ω from unity, corresponding to an interaction energy of 7.4 kJ/mol is relatively high ^33^, confirming that Q536 and D732 are energetically coupled. We tried to analyse energetic coupling between Q536 and A734 via double mutants incorporating nonsense suppression at the A734 position, but unfortunately, attempts to incorporate Aah into such double mutants via nonsense suppression were unsuccessful (Fig. S1B).

### L/F substitutions do not cause constitutive activity in all NMDA receptor subunits

We next asked if D732 substitutions have a similar effect on NMDA receptor activity when the substitution is present in other subunits than GluN1 or when mutant GluN1 subunits are combined with other WT subunits than GluN2A. GluN1/GluN3A receptors are glycine-gated channels that mediate relatively small currents in response to glycine, because despite glycine binding to and activating GluN3A subunits, glycine binding to GluN1 subunits deactivates the channel ^34,35^. GluN1/GluN3A receptors are assembled more efficiently by the GluN1(4a) isoform than the GluN1(1a) isoform that we used above ^36^, and therefore we used GluN1(4a) for our next experiments, after first confirming that the D732L substitution also generates solely glutamate-gated currents in GluN1(4a)-D732L/GluN2A-WT receptors (Fig. S3). Glycine alone activated no detectable currents in GluN1(4a)-D732L/GluN3A-WT receptors, and large glycine-gated currents were only observed in the presence of the GluN1-selective competitive antagonist CGP 78608, much like WT receptors (Fig. 5A) ^11^. While this seems like inactivity of the mutant GluN1 subunit in GluN1(4a)/GluN3A receptors, it could yet reflect the activity of constitutively active mutant GluN1 subunits in GluN1/GluN2A receptors, as GluN1 activity in GluN1/GluN3A receptors is known to cause more desensitization than activation ^11^. We also observed that the potency of glycine in activating GluN1(4a)/GluN3A receptors in the presence of CGP 78608 was decreased in GluN1(4a)-D732L-containing receptors, compared to WT (Fig 5B). Whether this derives from constitutive activity of GluN1(4a)-D732L inducing even greater channel deactivation than glycine binding to GluN1(4a)-WT, or from GluN1(4a)-D732L potentially having decreased CGP 78608 sensitivity, we did not explore.

**Figure 5.**
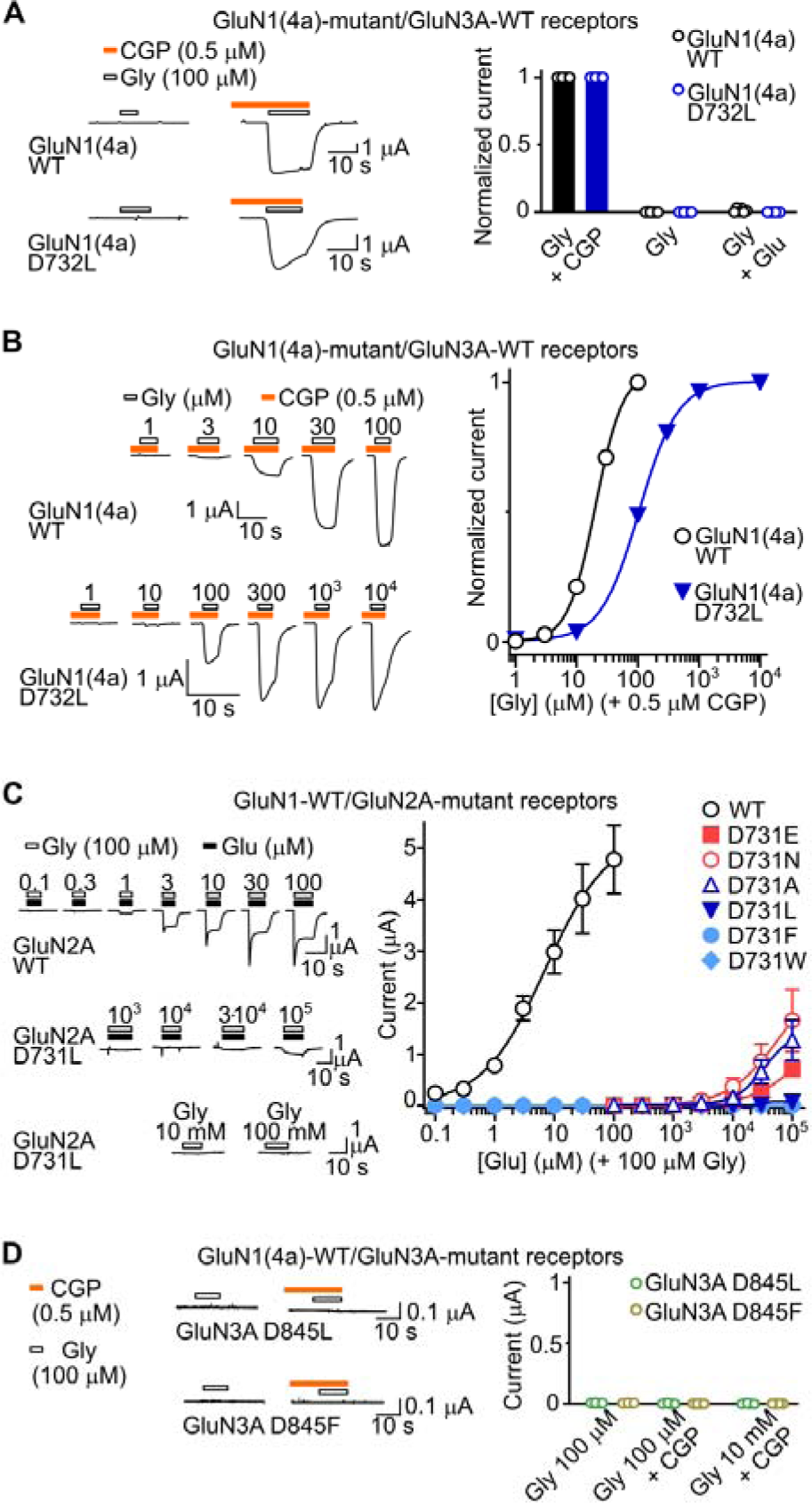
Effects of D732 substitutions in different NMDA receptor subtypes. (**A**) Left, example currents in oocytes expressing NMDA receptors comprising indicated WT or mutant GluN1 (4a isoform) subunits and WT GluN3A subunits in response to glycine alone or glycine in the presence of CGP 78608 (CGP). Right, Mean (columns) and individual data points (n = 4-6) for normalized current amplitude in the presence of different ligands. (**B**) Left, example currents in oocytes expressing NMDA receptors comprising indicated WT or mutant GluN1 (4a isoform) subunits and WT GluN3A subunits. Right, concentration-dependent responses (mean ± SEM, n = 6 (WT) or 5 (D732L)) for receptors carrying indicated GluN1 subunits. (**C**) Left, example ligand-gated currents in Xenopus laevis oocytes expressing NMDA receptors comprising WT GluN1 subunits and WT or mutant GluN2A subunits, as indicated. Right, concentration-current response relationships (data points mean ± SEM, n = 10 (WT), 3 (D731F and D731W), or 5-6 (all other mutants)) for receptors carrying WT GluN1 subunits together with indicated WT or mutant GluN2A subunits. (**D**) Example (left) and summarized (right) responses to indicated ligands in oocytes expressing WT GluN1 subunits (4a isoform) and indicated mutant GluN3A subunits.

Next, we measured the activity of receptors containing WT GluN1 subunits and mutant GluN2A or GluN3A subunits that contained substitutions equivalent to GluN1-D732L. GluN1-WT/GluN2A-D731L and -D731F receptors were not activated by glycine alone and were barely activated by high concentrations of glutamate in the presence of 100 μM glycine (Fig. 5C). We observed the same stark loss of function with the larger hydrophobic GluN2A-D731W substitution, and a less drastic but still >1000-fold decrease in glutamate potency with GluN2A-D731A/N/E substitutions (Fig. 5C). Similarly, at oocytes injected with GluN1(4a)-WT and GluN3A-D845L or -D845F mRNAs, we saw no response to glycine alone or glycine in the presence of CGP 78608 (Fig 5D). Together, these results indicate that among NMDA receptors the induction of a constitutively active state by the D732L/F substitution is an exclusive feature of the GluN1 subunit, and the only conserved feature of the mutation in different subunits is a decrease in agonist potency.

### Constitutive activity of a homomeric iGluR with a naturally occurring hydrophobic substitution

The effects of hydrophobic substitutions at the 732 position in the LBD thus differ among relatively complex NMDA receptors, in which the GluN1 subunits occupy the more accessory “A” and “C” positions in the hetero-tetramer (Fig. 1A, ^14^). We questioned if iGluRs of simpler homo-tetrameric architecture might be more susceptible to channel activation by similar hydrophobic substitutions. To answer this, we turned to the natural diversity of the iGluR superfamily and sought a wildtype receptor in which a naturally occurring substitution at the D732 (or equivalent) position might have been retained during evolution. We assembled diverse iGluR sequences, covering all animal lineages and animal iGluR families (Fig. 6A,B), and investigated variation at the D732 position. As expected, a carboxylate side chain at this position is broadly conserved, with glutamate or aspartate occurring here in 92% of the 204 iGluR genes in our analysis (Table 1). This level of conservation is higher than ligand-binding threonine/serine residue T518 (76%) and similar to pore-forming residue A653 (89%), but lower than ligand-binding R523 (97%) and pore-forming Y647 (99%, Table 1).

**Figure 6.**
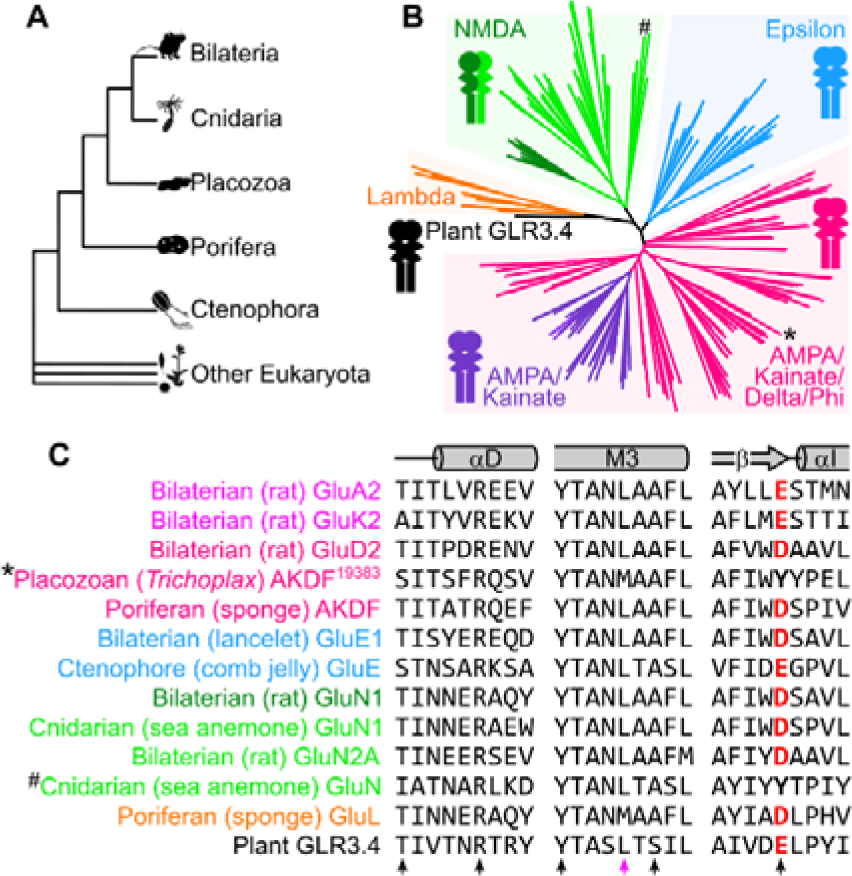
Conservation of D732 in diverse iGluRs. **(A)** Overview of major animal lineages. **(B)** Phylogeny of 204 animal iGluR genes, rooted to plant iGluR 3.4 outgroup (black tip), colored by phylogenetic families. Cartoons indicate obligate heteromeric (NMDA) or potential homomeric (others) iGluRs. Asterisk: Trichoplax AKDF^19383^ gene. Hash: a cnidarian GluN gene. **(C)** Amino acid sequence alignment of selected ligand-binding domain segments (helix αD and helix αI with preceding β-strand) and channel-forming segment (M3) from characterized (except for GluL and cnidarian GluN) members of each iGluR family. Position equivalent to GluN1 D732 highlighted red. Black arrowheads: residues addressed in Table 1. Pink arrowhead: residue addressed in Fig. 7D.

**Table 1.** Conservation of selected amino acid residues^a^.

|  | Amino acid residue (Rat GluN1 numbering) |  |  |  |  |
| --- | --- | --- | --- | --- | --- |
|  | T518 | R523 | Y647 | A653(Lc) | D732 |
| A | 7.8% | 0.0% | 0.0% | 88.7% | 0.5% |
| C | 0.0% | 0.0% | 0.0% | 0.0% | 0.0% |
| D | 0.0% | 0.0% | 0.0% | 0.0% | 54.9% |
| E | 0.0% | 0.0% | 0.0% | 0.0% | 37.3% |
| F | 0.5% | 0.0% | 1.5% | 0.0% | 0.0% |
| G | 2.0% | 0.5% | 0.0% | 0.5% | 3.4% |
| H | 0.0% | 0.5% | 0.0% | 0.5% | 1.5% |
| I | 3.4% | 0.5% | 0.0% | 0.0% | 0.0% |
| K | 3.4% | 0.5% | 0.0% | 0.0% | 0.0% |
| L | 2.5% | 1.5% | 0.0% | 0.0% | 0.0% |
| M | 0.5% | 0.0% | 0.0% | 0.0% | 0.0% |
| N | 0.0% | 1.5% | 0.0% | 0.0% | 0.0% |
| P | 1.5% | 0.0% | 0.0% | 0.5% | 1.0% |
| Q | 0.5% | 0.5% | 0.0% | 0.5% | 0.0% |
| R | 0.5% | 92.2% | 0.0% | 0.0% | 0.5% |
| S | 20.6% | 0.0% | 0.0% | 5.4% | 0.0% |
| T | 53.4% | 0.5% | 0.0% | 1.5% | 0.0% |
| V | 3.4% | 0.0% | 0.0% | 2.5% | 0.0% |
| W | 0.0% | 0.0% | 0.5% | 0.0% | 0.0% |
| Y | 0.0% | 2.0% | 98.0% | 0.0% | 1.0% |

<sup>a</sup> From alignment of 204 diverse iGluR subunits (Methods, Bioinformatic analyses).

Remarkably, in two genes we identified a large, hydrophobic tyrosine residue at the D732 position (Fig. 6B,C, ^#^ and *). One of these is v1g219279 from the sea anemone *Nematostella vectensis* ^37^, one of several genes forming a non-GluN1/2/3 cnidarian-specific group within the NMDA branch (Fig. 6C, ^5^). The other is *Trichoplax adhaerens* gene TRIADDRAFT_19383 (hence “AKDF^19383^”), from the broad AMPA/kainate/delta/phi (AKDF) branch of mostly homomeric iGluRs ^5^. Although not part of a characteristic group within the AKDF branch, such as AMPA or delta, AKDF^19383^ orthologues are conserved in distinct placozoans, including geographically distinct *Trichoplax* populations and also the more distantly related *Hoilungia hongkongensis* (Fig. S4A,B) ^38–40^. Moreover, each AKDF^19383^ orthologue has retained the tyrosine residue, despite divergence at other positions in the gene (Fig. S4A,B).

When we expressed *Trichoplax* AKDF^19383^ in *Xenopus* oocytes, we observed substantial current in the absence of ligands (390 ± 70 nA, n = 10 *cf.* 100 ± 20 nA, n = 8 in water-injected oocytes). The application of glycine, alanine, and D-serine but not glutamate or aspartate (each 10 mM) caused ∼50% *inhibition* of this current (Fig. 7A,B), indicating that AKDF^19383^ receptors are constitutively active, and ligand binding induces channel deactivation. To test if the constitutive activity is mediated by the Y742 residue, we tested the activity of mutant AKDF^19383^ receptors in which Y742 is replaced with a more typical aspartate residue (Y742D). Remarkably, glycine and D-serine, and to a lesser extent glutamate, activated inward currents through Y742D receptors (Fig. 7A, mean glycine-gated iwnard current 1.8 ± 0.6 μA, n = 7). Ligand-gated currents were rapidly activating and deactivating in the continued presence of ligands (Fig. 7A,C), reflecting “desensitization” common to several iGluRs of the AKDF family, such as mammalian AMPA and kainate receptors ^4^. Constitutive currents were still present in Y742D receptors but in contrast to WT, these were not inhibited by the pore blocker pentamidine or by most agonists (Fig. 7A,B). The agonist glycine, however, induced desensitization that exceeded glycine-gated inward current amplitude, such that glycine effectively inhibited some of the constitutive current in Y742D receptors (Fig. 7C). Thus, a tyrosine or aspartate residue at the 742 position determines whether AKDF^19383^ is a constitutively active receptor inhibited by ligand-binding and conventional pore-blockers or is instead a principally ligand-gated channel that retains some constitutive activity.

**Figure 7.**
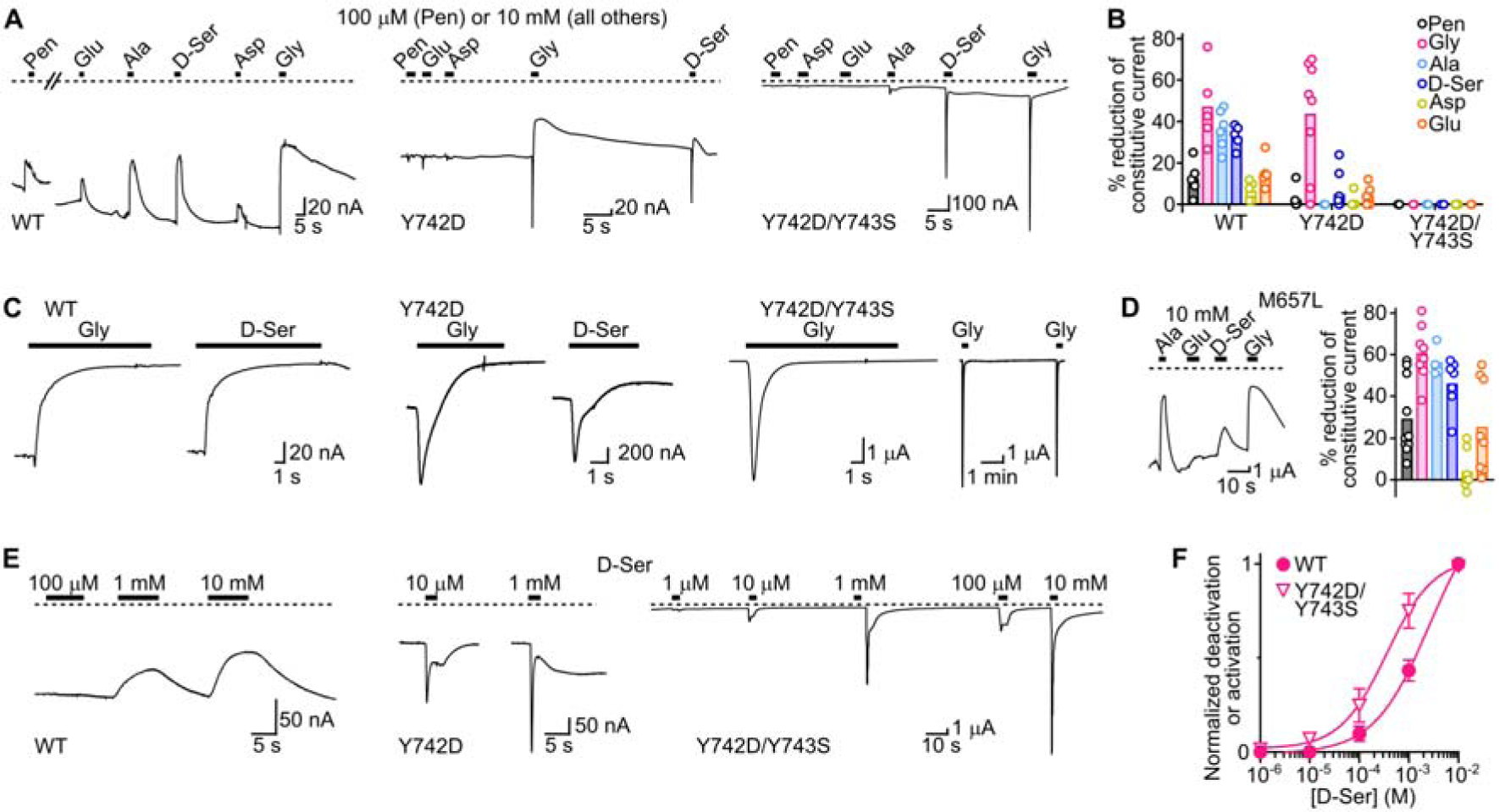
Function of Trichoplax AKDF^19383^ iGluR. **(A)** Constitutive current (dashed line = zero current) and responses to different ligands in oocytes expressing indicated wildtype (WT) or mutant Trichoplax AKDF^19383^ receptors. Asp, L-aspartate. Glu, L-glutamate. D-Ser, D-Serine. Gly, glycine. Ala, alanine. Pent, pentamidine, an iGluR channel blocker. **(B)** Summary data from (**A**): individual data points and means (columns) are shown. **(C)** Magnified views of glycine and D-serine activity from (**A**) (longer horizontal bars), and additional recording (far-right) showing decreasing amplitude of responses to glycine at Y742D/Y743S receptors. **(D)** Example recording (left, dashed line = zero current) and summary data (columns = means, n = 5-9) showing inhibition of constitutive currents in oocytes expressing mutant M657L Trichoplax AKDF^19383^ receptors. Colors as in (**B**). **(E)** Current responses to increasing concentrations of D-serine in oocytes expressing indicated Trichoplax AKDF^19383^ receptors. **(F)** Mean ± SEM (n = 4) deactivation normalized to maximum deactivation for WT and activation normalized to maximum activation for mutant Y742D/Y743S Trichoplax AKDF^19383^ receptors.

We noticed that *Trichoplax adhaerens* AKDF^19383^ has an additional tyrosine residue immediately following Y742, again differing from most other iGluRs – including the *Holiungia hongkongensis* orthologue – which instead typically possess a small serine, glycine, or alanine at this position (Fig. 6C, Fig. S4A). We therefore generated and tested double-mutant Y742D/Y743S AKDF^19383^ receptors and now observed large ligand-gated currents and a nearly total reduction of the constitutive current (Fig. 7A,C). This shows that amino acid residues at LBD positions 742 and 743 together determine the channel activity of the *Trichoplax* AKDF^19383^ iGluR. As channel activity of iGluRs also relies on the upper segment of the third membrane-spanning helix (M3, ref. ^41^), we also examined this segment in AKDF^19383^. AKDF^19383^ differs only subtly from most iGluRs here, with a methionine residue (M657) instead of a typical leucine (Fig. 6C, pink arrowhead), but we tested potential effects of this small difference by mutating M657 to leucine. Despite apparently larger leak current amplitude, M657L activity was much like WT (Fig. 7D), confirming that divergence at Y742/Y743, and not the upper M3 segment, determines the unique activity of AKDF^19383^.

Finally, we tried to assess whether position 742 determines ligand potency in addition to channel activity in AKDF^19383^ receptors. For these experiments we employed D-serine, as recovery from glycine-induced deactivation (Fig. 7C, far-left) and activation/desensitization (Fig. 7C, far-right) was slow. Substantial deactivation of WT receptors was only induced by millimolar D-serine concentrations, whereas Y742D-containing mutants were activated by micromolar concentrations (Fig. 7E,F), with an EC_50_ of 490 ± 120 µM at Y742D/Y743S (n = 4; Y742D EC_50_ not assessed due to slow recovery from desensitization). Our measure of potency is confounded by the fact that deactivation (in WT channels) and activation (in mutant channels) are presumably coupled to D-serine binding via different conformational transitions. Nonetheless, we observe that a naturally occurring large hydrophobic side chain at the top of the β-strand preceding the αI helix leads to an AKDF homo-tetramer that shows constitutive activity and responds only to millimolar concentrations of D-serine. In contrast, “re-introducing” an aspartate to this position reinstates more typical ligand-dependent activation and sensitivity to micromolar concentrations of D-serine. This is remarkably consistent with results in the GluN1 subunit of distantly related NMDA receptors, pointing toward a dominant role of this residue in LBD activation throughout the iGluR superfamily.

## DISCUSSION

Seeking to understand which amino acid residues control iGluR activation and to facilitate the prediction of function from sequence, we investigated the role of a broadly conserved carboxylate residue in the middle of the iGluR LBD. Our experiments with heteromeric NMDA receptors and with a distantly related homomeric AKDF iGluR suggest that this residue controls agonist potency in most iGluRs and has a profound link to channel gating in certain iGluRs.

### D732 in ligand binding

That D732 should affect agonist potency in diverse iGluRs is not surprising. Previous reports show decreased agonist potency in NMDA receptors and AMPA receptors carrying mutations at the homologous aspartate and glutamate residue, respectively ^24,42^, and X-ray structures of isolated GluN1, GluN2A, and GluA2 LBDs place the carboxylate ∼3 Å from the amine of agonists glycine or glutamate ^15,16,43^. Our study suggests that D732 conformational change, and simply amino acid identity in unliganded receptors, controls channel activity, warranting further consideration of this residue in future studies dissecting the conformational changes that underlie iGluR activation.

Several ligand-induced conformational changes occur during ligand binding and before channel opening, as shown by kinetic descriptions of NMDA receptor single channel openings and closures (e.g. refs.^44,45^). Distinct pre-gating steps have also been identified in AMPA receptors via time-resolved fluorescence- and infrared-based analyses: a fast, preliminary “docking” of ligand to the upper lobe; and then a slower “locking” in which the lower lobe closes around the ligand ^46,47^. And remarkably, E726 (≈ GluN1 D732) mutations primarily impair the slower, locking reaction ^46,47^. If these pre-gating steps are similar in NMDA receptors, it is easy to imagine that glycine/glutamate binds to GluN1/GluN2 upper lobes during docking, and that the 732 side chain swings toward the ligand amine during locking. Indeed, in a cryo-electron microscopy (cryo-EM) study of GluN1/GluN2B receptors, D732 has only swung toward the ligand and away from A734 in a second of two putative pre-gating step structural models, although this is speculative considering the poor resolution of D732 side chains in those cryo-EM maps ^14^. If the D732 side chain is shorter, longer, and or non-carboxylate, locking presumably becomes less favourable, potentially explaining the decreased agonist potency at GluN1-D732A/S/N/E/Y/W, GluN2A-D731A/N/E/L/F/W, and GluN3A-D785L/F mutants, and at AKDF^19383^ WT receptors.

### D732 or hydrophobic substituents in channel activation

Higher resolution X-ray structures of isolated NMDA receptor LBDs capture the D732 carboxylate in two distinct conformations, implying the side chain swings ∼3 Å toward glycine during activation (Fig. 4A, ^15,23,48^). Further analysis of these structures shows D732 movement is intricately coupled to Q536 side chain reorientation, which is supported by our experiments suggesting that interactions between hinge region-Q536, helix αI-A734, and the agonist contribute to glycine potency in the GluN1 LBD. Thus, D732 could contribute to locking and further conformational changes by mediating coupling between (a) glycine binding and (b) hinge and αI conformation. Similar coupling was suggested for AMPA receptors, where ligand binding links the rearrangement of E726 during a late ligand-binding step to substantial lower lobe movement and even secondary structure changes, especially in the β-sheet below E726 in AMPA receptors ^46,49,50^. How this LBD movement is then transduced to channel opening is beyond our study, but as the LBD-M1, -M3, and -M4 linkers play distinct roles in later pre-gating steps ^13,14^, perhaps future studies may link D732 rearrangement to distinct structural changes closer to the channel.

GluN1 and AKDF^19383^ subunits with a certain hydrophobic substituent at position 732 (GluN1 numbering) apparently overcome most of these pre-gating and gating steps in the absence of ligand. As glycine-free mutant and glycine-bound WT NMDA receptors show similarities in apparent open probability and redox sensitivity, it’s tempting to conclude that the hydrophobic substituent closes the mutant LBD the same way glycine closes WT LBDs. Perhaps the hydrophobic side chain re-positions water molecules linked to ligand-induced LBD closure ^51^, bridges the lower lobe directly to upper lobe hydrophobic side chains, and/or allows better access of β12-Q536 to αI-A734. However, we cannot rule out subunit activation by hydrophobic side chains without LBD closure, as the mutant GluN1 subunits are potently inhibited by the large antagonist DCKA, and published GluN1 X-ray structures show less agonist-dependent LBD closure than other iGluRs ^21^.

Another possibility is that the hydrophobic side chain-containing LBD cooperates differently with adjacent subunits. Indeed, glycine activity at GluN1 subunits negatively impacts glutamate- and glycine-induced activation of GluN2 and GluN3A subunits, respectively ^11,52^, suggesting that simply reducing glycine activity could enhance NMDA receptor activation by glutamate. However, most GluN1 mutations that decreased glycine sensitivity led to no glutamate-gated currents here, and DCKA binding to GluN1 inhibited channel activation by glutamate. Curiously, positive cooperativity between GluN1 and GluN2 subunits has been observed for certain combinations of ligands ^53^, underscoring that inter-subunit interactions are complex and deserve more investigation. Molecular dynamics simulations correlating amino acid rearrangements with LBD closure ^23,51^ and structural studies of mutant GluN1-D732L/F NMDA receptors or WT AKDF^19383^ receptors could foreseeably answer the precise mechanisms by which hydrophobic substituents induce channel activation.

### Physiological implications

iGluRs mediate most excitatory signals in the mammalian brain, and whereas AMPA receptors do the bulk of this work, NMDA receptors make vital contributions because of their distinct functional properties. These include high calcium ion permeability, the requirement for prior depolarization, and via their obligate GluN1/GluN2 heteromeric nature the requirement for two different ligands.

Our work shows that only small changes in the GluN1 LBD are required for solely glutamate-gated currents in vertebrate GluN1/GluN2 receptors, and previous work suggests that invertebrate *Drosophila melanogaster* and *Aplysia californica* GluN1/GluN2 receptors can be activated by single ligands ^54,55^. This suggests that NMDA receptors’ requirement of co-agonism is easily alleviated by certain mutations or conditions. As iGluR-modulatory proteins vary across cell types or even across neuronal compartments ^56,57^ and NMDA receptor sequence varies across animals, it is foreseeable that in certain physiological settings, certain NMDA receptors might be activated by glutamate alone. But in most settings, certainly in vertebrates, it seems that glutamate-induced activation of NMDA receptors relies on a system of ambient glycine or D-serine ^58,59^.

Genetic knock-in of solely glutamate-gated NMDA receptors via a single D732L/F mutation would allow the comparison of co-agonist-dependent and -independent NMDA receptors on other mutant backgrounds (e.g. phgdh^-/-^ or SR^-/-^ ^59,60^) to establish the precise contributions of co-agonists to NMDA receptor activity *in vivo*. Solely glutamate-gated and glycine-gated mutant rat NMDA receptors were reported previously after the introduction of pairs of cysteine residues in the upper and lower lobes of GluN1 or GluN2A subunits, respectively ^26,27^. But as these rely on redox environment, which modulates WT NMDA receptors via native GluN1 disulfides ^61^, these cysteine-mutants are less suited to knock-in studies.

### Biological implications

The dominant role of a broadly conserved residue in agonist potency and channel activity prompted us to ask if we could predict function from amino acid sequence. Predicting *ligand potency* based on amino acid identity at the 732 (or equivalent) position holds for GluN1, GluN2A, and GluN3 NMDA receptor subunits as well as *Trichoplax* AKDF^19383^ receptors, as an aspartate residue is conducive to higher potency and other residues to lower potency. But predicting *channel activity* – ligand-gated or constitutively active – simply based on identity of this residue is more tenuous, as among NMDA receptors the induction of a constitutively active subunit by the L/F substitution is unique to GluN1 subunits. It is remarkable, however, that even in the distantly related homotetrameric AKDF^19383^ receptor, we see a similar pattern to GluN1, where an aspartate residue at this position is conducive to potently ligand-gated currents, and a large hydrophobic tyrosine side chain causes decreased ligand potency and constitutive channel activity. The pattern is not absolutely conserved, however, as only leucine and phenylalanine – not tyrosine – caused constitutive activity of GluN1 subunits.

The identification of a constitutively active iGluR channel from placozoans is striking. This adds iGluRs to the list of ligand-gated channel superfamilies in which leak channels have been identified, also including for example degenerin/epithelial sodium channels and pentameric ligand-gated ion channels ^62,63^. AKDF^19383^ orthologues are conserved in all three identified strains of placozoans, simple, flat animals with relatively few cell types ^64^. Single-cell RNA sequencing of *Trichoplax adhaerens* suggests that AKDF^19383^ is expressed in lipophil cells ^65^. These are large, unciliated columnar cells, interspersed between ventral epithelial cells and essentially bridging the ventral exterior with the interior of the animal ^66^. Classical neurons or synapses are probably absent from placozoans, as inferred from co-expression of typical pre- and post-synaptic genes in *Trichoplax adhaerens* ^40,65^, but previous reports of altered feeding in response to bath application of glycine and glutamate ^67^ and intercellular peptidergic signals from ventral ciliated cells ^68^ are indicative of chemosensory and integrative functions in these animals. Whether the AKDF^19383^ receptor behaves the same way in living placozoans as in our heterologous experiments is unknown, but we envisage it as a leak channel that is inhibited by small amino acid ligands either released from adjacent cells or found on the animal’s substrate. Consideration of the equivalent hydrophobic/carboxylate residue in future sequencing projects will likely help distinguish constitutively active from typical, ligand-gated receptors of the iGluR superfamily.

## METHODS

### Plasmids and molecular biology

*Rattus norvegicus* GluN1 (1a isoform) and GluN2A (the regular, long isoform) inserts within the multiple cloning site (MCS) of pcDNA3.1 without poly(A) tails were a gift from Dr. David MacLean (University of Rochester Medical Center). *Rattus norvegicus* GluN1 (4a isoform) and GluN3A containing a C-terminal DYKDDDDK and within the MCS of pcDNA3.1 were purchased from Genscript (ORa13617 and ORa44363, respectively). The *Trichoplax adhaerens* AKDF^19383^ sequence was established by additional genome annotation of the Genbank TRIADDRAFT_19383 sequence (described in Fig. S4 legend), codon-optimized for *Xenopus laevis* in iCodon ^69^, and commercially synthesized and cloned into a modified pSP64-poly(A) vector (full sequence in Supplemental text).

Mutations were generated with partially overlapping primers and PCR with Phusion Hot Start II DNA Polymerase (ThermoFisher) according to published guidelines ^70^. Inserts were fully sequenced via Sanger sequencing.

Plasmids were linearized via restriction digest at sites downstream of the coding sequence (Rat GluN1/2/3) or downstream of the poly(A) segment (*Trichoplax* AKDF^19383^) and purified via ethanol precipitation or DNA Clean & Concentrator-5 columns (Zymo Research). mRNA was transcribed for 2-2.5 h with mMESSAGE mMACHINE kits (ThermoFisher, T7 for pcDNA3, SP6 for pSP64-poly(A)) following supplier’s protocols including DNAse treatment and purified with RNA Clean & Concentrator-5 (Zymo Research) or RNAeasy (Qiagen) kits.

### Non-canonical amino acid chemistry

#### General

All reactions were carried out using commercial materials and reagents without further purification unless otherwise noted. All reactions were monitored by thin layer chromatography (TLC) on silica gel plates (Keiselgel 60 F254, Merck) and/or ultra-performance liquid chromatography (UPLC). Visualization of the spots on TLC plates was achieved by UV light and by staining the TLC plates in potassium permanganate and charring with a heat gun, unless otherwise stated.

UPLC refers to a Waters Acquity UPLC HClass instrument with Acquity PDA detector, QDA mass detector and quaternary solvent system; PDA: 210-350 nm. Acidic methods were run using varying gradients of acetonitrile and water with 5% 2 vol% formic acid (99%) in water on the following columns: Acquity CSH C18 column (2.1 x 50 mm 1.7 µm) at 0.8 mL/min; Acquity CSH-Phenyl Hexyl (CSH PH) column (2.1 x 50 mm 1.7 µm); HSS T3 column (2.1 x 50 mm 1.8 µm) at 0.8 mL/min. Basic methods were run using varying gradients of acetonitrile and water with 5% 2 vol% ammonia (28%) in water on the following columns: XBridge BEH C18 column (2.5 µm 2.1 x 50 mm) at 0.8 mL/min; XBridge BEH C8 column (2.5 µm 2.1 x 50 mm) at 0.8 mL/min. Eluents A: water, B: acetonitrile, C: 2 vol % ammonia (28%) in water, D: 2 vol % formic acid (99%) in water.

Products characterized by ^1^H NMR was recorded on a Bruker Advance III HD 400 MHz spectrometer. Chemical shifts are expressed in parts per million values (ppm) and are designated as s (singlet); br s (broad singlet); d (doublet); t (triplet); q (quartet); quint (quintet) or m (multiplet).

Chromatography was performed on a Biotage Isolera using silica (normal phase) (SiliCycle SiliaSep Premium 25 µm or Biotage SNAP Ultra HP-Sphere 25 µm) or C18 (reverse phase) (Biotage SNAP Ultra C18 HP Sphere 25 µm) pre-packed cartridges; or by flash-column chromatography using silica gel (Fluorochem silica gel 60A 40-63 µm).

**General procedure A**

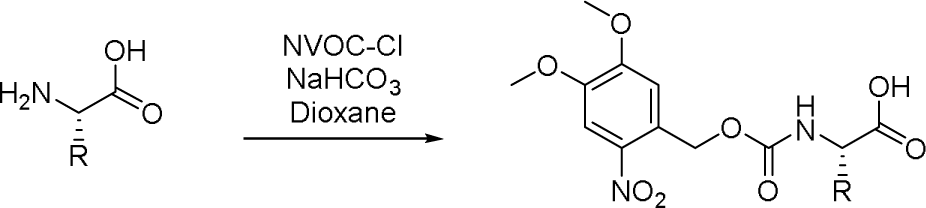

A suspension of amino acid (1 eq), (4,5-dimethoxy-2-nitro-phenyl)methyl carbonochloridate (1.1 eq) and sodium hydrogen carbonate (2.5 eq) in 1,4-dioxane (106 eq) was stirred at room temperature overnight. Water was added and stirred at room temp for 30 min. The reaction was diluted with ethyl acetate and the phases separated. The organic phase was discarded. The aqueous was acidified to ∼pH 1 with 2 M HCl aqueous solution. The product was extracted with ethyl acetate and dried over MgSO_4_. The solution was concentrated under vacuum to give the desired product which was used without purification in the next step.

**General procedure B**

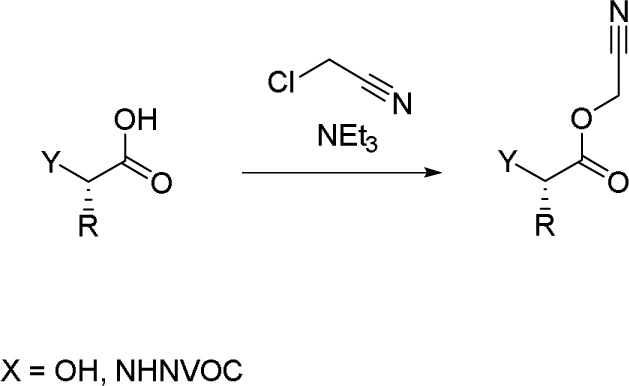

A stirring solution of NVOC-amino acid or 2-subsituted-2-hydroxyacetic acid (1 eq) and triethylamine (2 eq) in chloroacetonitrile (50 eq) was stirred at room temp overnight. The reaction mixture was concentrated under vacuum. The residue was suspended in EtOAc and sonicated for 10 min. The mixture was filtered and the filtrate concentrated under vacuum to give the desired cyanomethyl ester. Used in the next step without purification.

**General procedure C**

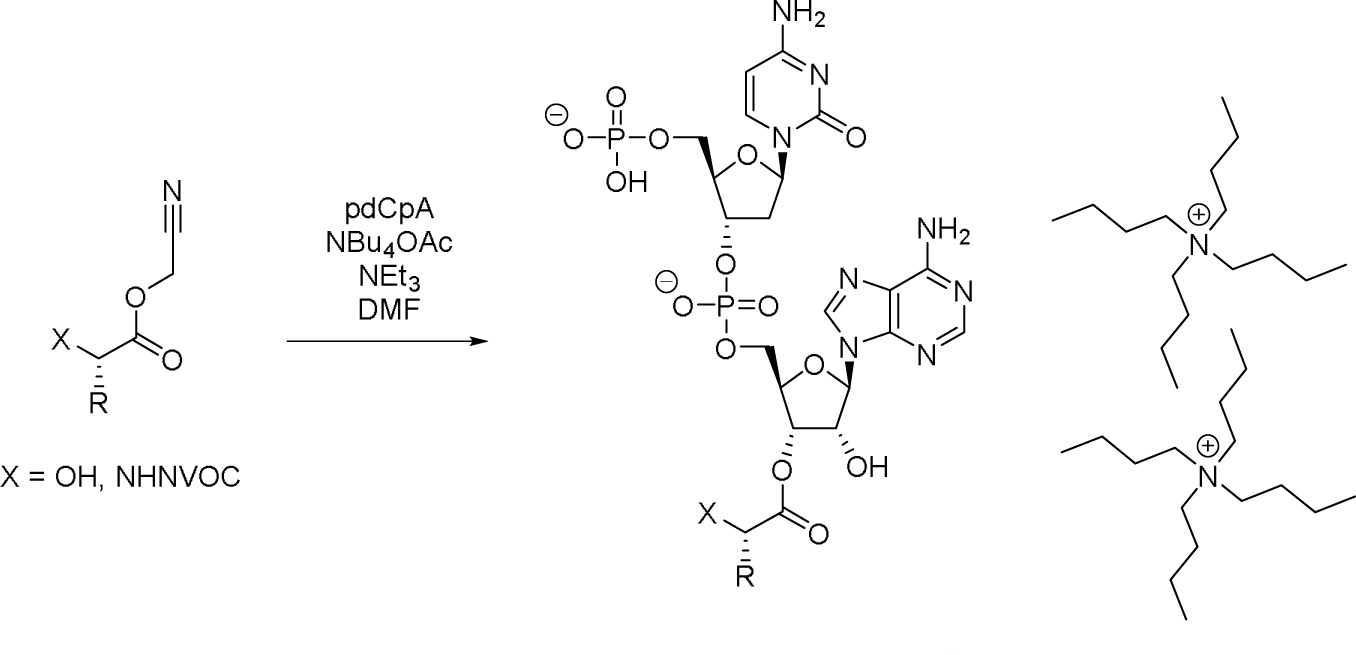

A solution of pdCpA (1 eq, synthesized according to ^71^), cyanomethyl ester (1 eq), tetrabutylammonium acetate (2 eq) and triethylamine (2 eq) in *N*,*N*-dimethylformamide (70 eq) was stirred at 38 °C overnight. The reaction mixture was directly purified by reverse phase purification (Biotage Isolera, 12 g SiliaSep C18 cartridge; gradient 20-70% (acetonitrile + 0.1%formic acid) in (water + 0.1% formic acid) over 12 CV). The product containing fractions were freeze dried overnight to give the desired compound as a mixture of monoacylated isomers at the 3’ and 2’ positions.

**Ditetrabutylammonium Nvoc-Val-OpdCpA**

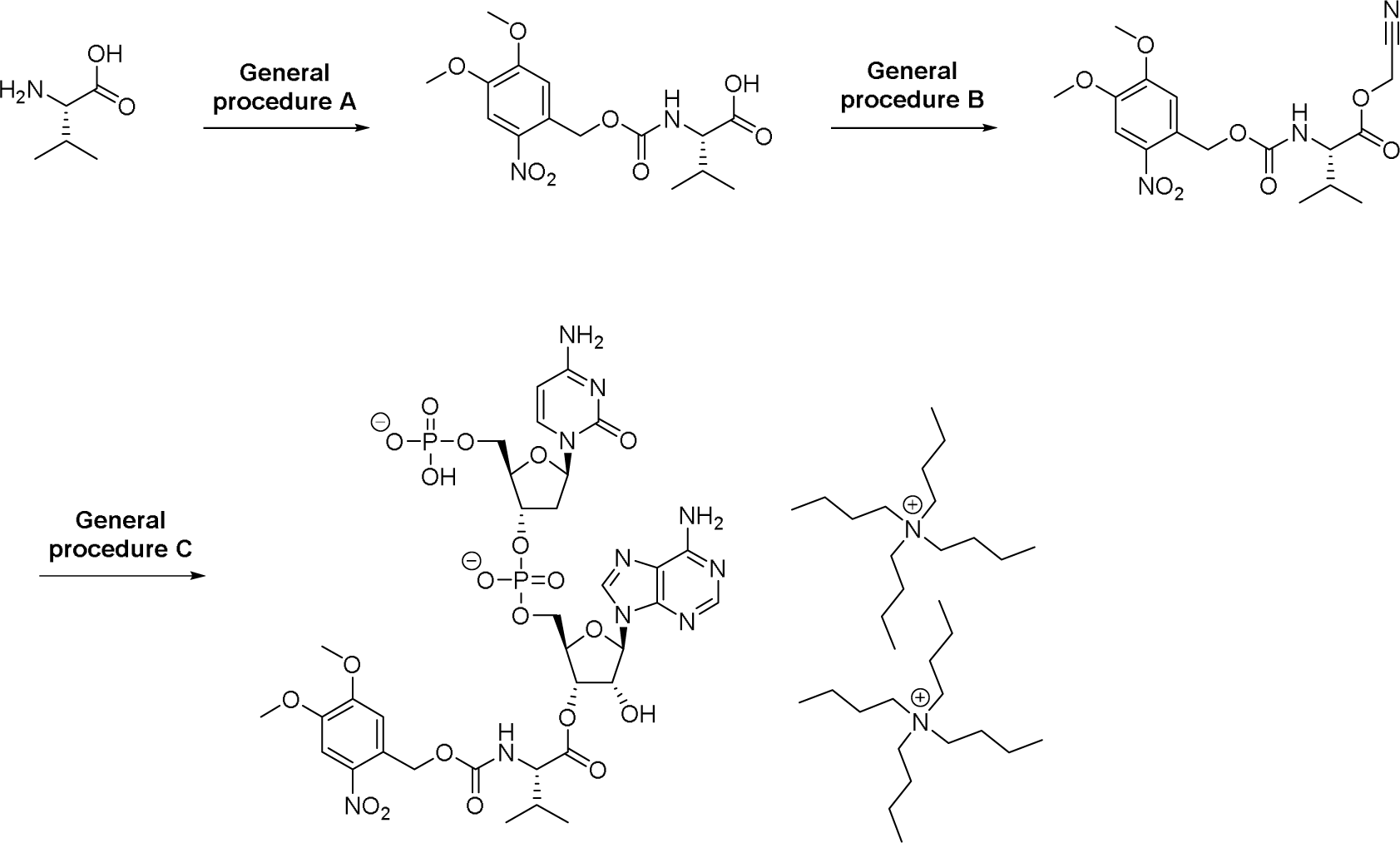

Using general procedure A, *L*-valine (58.0 mg, 1 eq, 0.495 mmol) gave the desired product Nvoc-Val-OH as yellow solid, 130 mg (74%). It was used in the next step without purification. UPLC-MS: (CSH-C18 Short Acid 2 to 95%, 1204-044-1) R_t_ = 0.79 min (83.7%), MS (ESIneg): m/z = [M-H]^-^ 355.

Using general procedure B, Nvoc-Val-OH (119 mg, 1 eq, 0.334 mmol) gave the desired product Nvoc-Val-OCH_2_CN as a yellow solid, 129 mg (98%). It was used in the next step without purification. UPLC-MS: (CSH-C18 Short base 2 to 95%, 1204-050-1) R_t_ = 0.86 min (90.7%), MS (ESIpos): m/z = [M+Na]^+^418.

Using general procedure C, Nvoc-Val-OCH_2_CN (12.0 mg, 1 eq, 31.4 μmol) gave the desired product, ditetrabutylammonium Nvoc-Val-OpdCpA as a white solid, 4.52 mg (9.3%).

UPLC-MS: (HSS T3 Long acid 0 to 10%, NBK2001-115-P1-2C) R_t_ = 1.78 min (30.2%), MS (ESIpos): m/z = [M+H]^+^ 975; MS (ESIneg): m/z = [M-H]^-^ 973 and 1.82 min (64.4%), MS (ESIpos): m/z = [M+H]^+^ 975; MS (ESIneg): m/z = [M-H]^-^ 973 – Combined purity of 94.6% (mixture of monoacylated isomers) (Fig. S5).

**Ditetrabutylammonium Vah-OpdCpA**

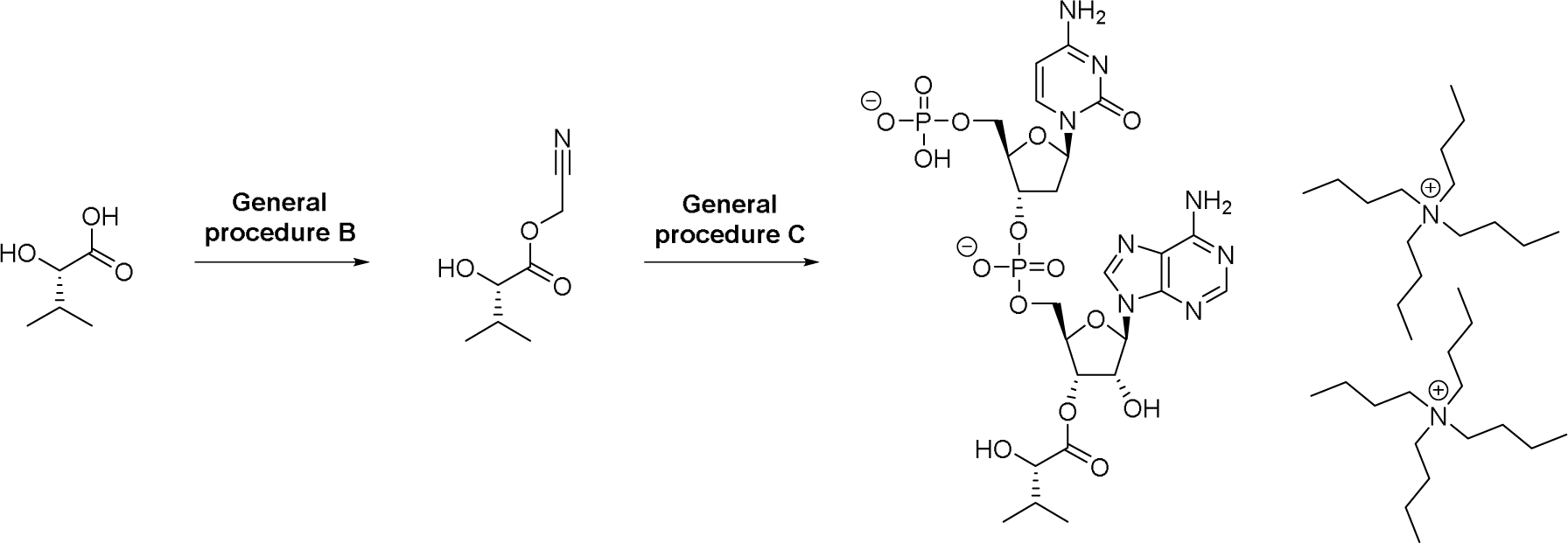

Using general procedure B, (*S*)-2-hydroxy-3-methylbutanoic acid (100 mg, 1 eq, 0.847 mmol) gave the desired product Vah-OCH_2_CN as a colorless oil, 110 mg (83%). It was used in the next step without purification. ^1^H NMR (400 MHz, CDCl_3_, NBK2001-112-P1-1): δ [ppm] = 0.91 (d, 3H), 1.04 (d, 3H), 2.08-2.16 (m, 1H), 4.15 (d, 1H), 4.82 (q, 2H). 1 exchangeable proton not observed.

Using general procedure C, Vah-OCH_2_CN (4.94 mg, 1 eq, 31.4 μmol) gave the desired product, ditetrabutylammonium Vah-OpdCpA as a white solid, 0.79 mg (2%).

UPLC-MS: (HSS T3 Long acid 0 to 10%, NBK2001-114-P1-2) R_t_ = 2.26 min (19.5%), MS (ESIpos): m/z = [M+H]^+^ 737; MS (ESIneg): m/z = [M-H]^-^ 735 and 2.60 min (80.1%), MS (ESIpos): m/z = [M+H]^+^ 737; MS (ESIneg): m/z = [M-H]-735 – Combined purity of 99.6% (mixture of monoacylated isomers). (Fig. S6.)

**Ditetrabutylammonium Nvoc-Ala-OpdCpA**

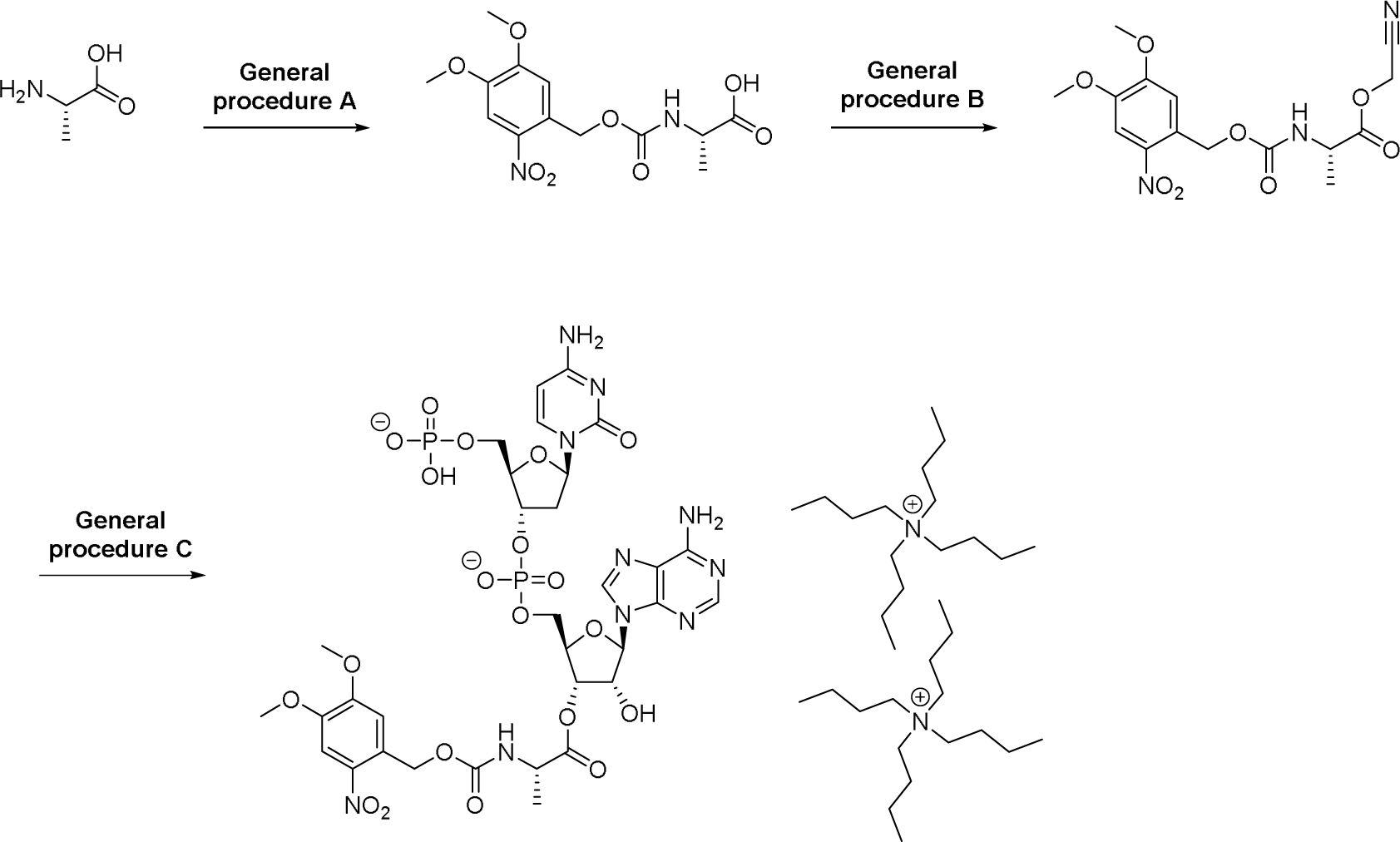

Using general procedure A, *L*-alanine (44.1 mg, 1 eq, 0.495 mmol) gave the desired product Nvoc-Ala-OH as yellow solid, 119 mg (73%). It was used in the next step without purification. UPLC-MS: (CSH-C18 Short Acid 2 to 95%, 1204-045-1) R_t_ = 0.68 min (84.1%), MS (ESIpos): m/z = [M+Na]^+^ 351; MS (ESIneg): m/z = [M-H]^-^ 327.

Using general procedure B, Nvoc-Ala-OH (119 mg, 1 eq, 0.362 mmol) gave the desired product Nvoc-Ala-OCH_2_CN as a yellow solid, 128 mg (96%). It was used in the next step without purification. UPLC-MS: (CSH-C18 Short Acid 2 to 95%, 1204-051-1) R_t_ = 0.75 min (78.8%), MS (ESIpos): m/z = [M+Na]^+^ 390.

Using general procedure C, Nvoc-Ala-OCH_2_CN (30.0 mg, 1 eq, 47.1 μmol) gave the desired product, ditetrabutylammonium Nvoc-Ala-OpdCpA as a white solid, 17.0 mg (38%).

UPLC-MS: (HSS T3 Long acid 0 to 20%, 1273-062-2) R_t_ = 2.85 min (46.6% at 254 nm), MS (ESIpos): m/z= [M+H]^+^ 947; MS (ESIneg): m/z = [M-H]^-^ 945 and 3.03 min (48.8% at 254 nm), MS (ESIpos): m/z = [M+H]^+^ 947; MS (ESIneg): m/z = [M-H]^-^ 945 – Combined purity of 95.4% (mixture of monoacylated isomers). (Fig. S7.)

**Ditetrabutylammonium Aah-OpdCpA**

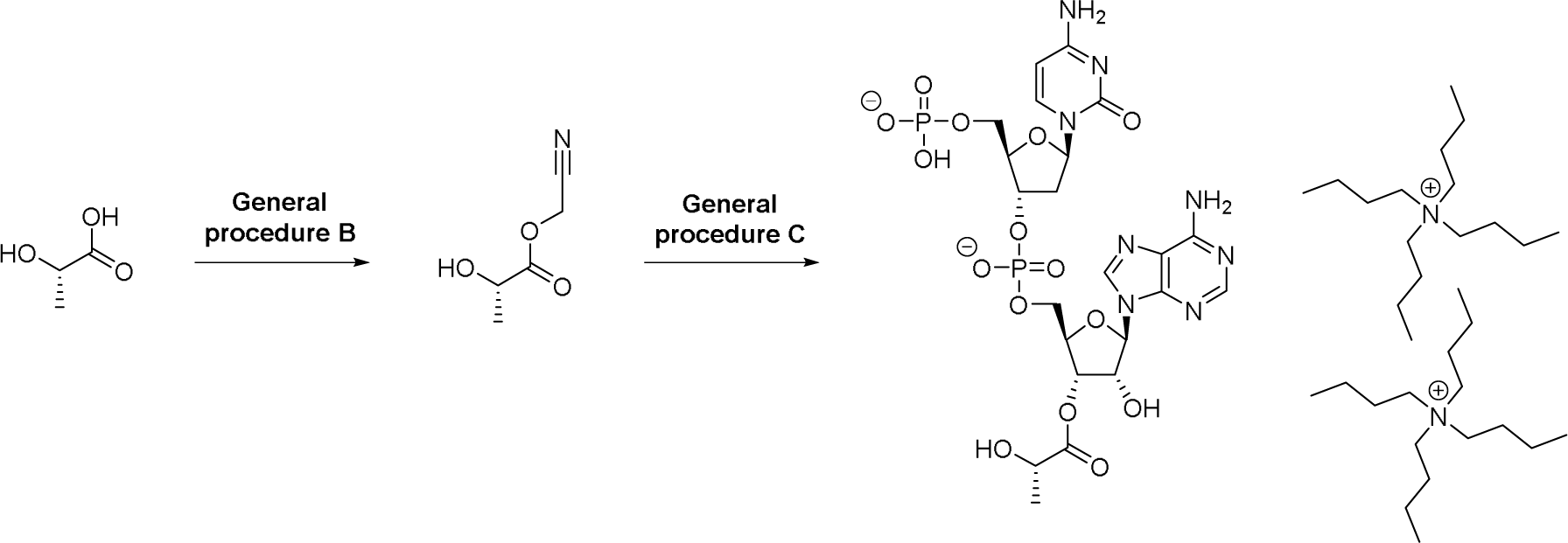

Using general procedure B, (*S*)-2-hydroxypropanoic acid (100 mg, 1 eq, 1.11 mmol) gave the desired product Aah-OCH_2_CN, 95.0 mg (66%) as a colorless oil. It was used in the next step without purification. ^1^H NMR (400 MHz, CDCl_3_, NBK2001-111-P1-1): δ [ppm] = 1.47 (d, 3H), 4.41 (q, 1H), 4.81 (s, 2H). 1 exchangeable proton not observed.

Using general procedure C, Aah-OCH_2_CN (4.06 mg, 1 eq, 31.4 μmol) gave the desired product, ditetrabutylammonium Aah-OpdCpA was isolated as a white solid, 2.60 mg (6.4%).

UPLC-MS: (HSS T3 Long acid 0 to 10%, NBK2001-113-P1-2) Rt = 1.55 min (22.3%), MS (ESIneg): m/z = [M-H]^-^ 707 and 1.75 min (69.9%), MS (ESIpos): m/z = [M+H]^+^ 709; MS (ESIneg): m/z = [M-H]^-^ 707 – Combined purity of 92.2% (mixture of monoacylatedisomers). (Fig. S8.)

### Channel expression and non-canonical amino acid incorporation

Defoliculated Stage V/VI *Xenopus laevis* frog eggs were purchased weekly from EcoCyte Bioscience and stored at 18 °C in 50% (in water) Leibovitz’s L-15 medium (Gibco) supplemented with additional 0.25 mg/ml gentamicin, 1 mM L-glutamine, and 15 mM HEPES, pH 7.6.

For the expression of wildtype and conventional mutant channels, oocytes were injected with 2 ng (rat NMDA receptors) or 10 ng (*Trichoplax* AKDF^19383^) RNA in 40 nl water using a Nanoliter2010 injector and glass micropipettes (World Precision Instruments). Non-canonical amino acid mutant NMDA receptor channels were expressed by co-injecting oocytes with amber stop codon mutant GluN1(1a) mRNA, wildtype GluN2A mRNA, and aminoacylated tRNA ^72^. Modified *Tetrahymena thermophila* tRNA^Gln^ ^73^ was prepared by ligation of forward and reverse DNA strands containing the T7 promotor (Merck), transcription of RNA with the T7-Scribe transcription kit (Cellscript), and purification in Chroma Spin DEPC-H20 columns (Clontech). This tRNA was ligated to Nvoc-Val-OpdCpA, Nvoc-Ala-OpdCpA, Vah-OpdCpA, and Aah-OpdCpA with T4 DNA ligase (New England Biolabs), and the product was purified with phenol-chloroform extraction and ethanol precipitation and stored at −80°C until use. Immediately before injection into oocytes, precipitate was resuspended in 1 µl water, deprotected via UV light (Nvoc-Val-OpdCpA and Nvoc-Ala-OpdCpA only) in a coil of 365 nm realUV LED strip lights (Waveform Lighting) cooled with a fan, and mixed 2:1:1 with 800 ng/μl amber stop codon mutant GluN1 mRNA and 200 ng/μl wildtype GluN2A mRNA (Fig. S1). 40 nl of this mix was then injected into oocytes. Injected oocytes were stored in the same diluted L-15 medium (above) until electrophysiological recordings.

### Two-electrode voltage clamp and data analysis

One to two days (rat NMDA receptors) or three to four days (*Trichoplax* AKDF^19383^) after injection, each oocyte was placed in an RC-3Z bath (Warner Instruments) and continuously perfused with a bath solution of NaCl 100 mM, KCl 2.5 mM, BaCl_2_ 0.3 mM, HEPES 5 mM, diethylenetriamine-pentaacetic acid 0.01 mM, pH 7.3 (rat NMDA receptors, to avoid inhibition by magnesium and zinc ions ^74^) or NaCl 96 mM, KCl 2 mM, CaCl 1.8 mM, MgCl 1 mM, HEPES 5 mM, pH 7.5 (*Trichoplax* AKDF^19383^). Bath solution was exchanged for that containing ligand as indicated in figures with a VCS-8-pinch valve control perfusion system (Warner Instruments). Ligands were stored as 1 M stocks in water (glycine) or 10 – 300 mM stocks in water or bath solution pH adjusted with NaOH (CGP 78608, monosodium glutamate, DCKA) and dissolved in bath solution before use. CGP 78608 was purchased from Tocris, all others from Merck. Oocytes were impaled with silver-chloride microelectrodes in 3M KCl-filled borosilicate micropipettes and clamped at −60 mV (rat NMDA receptors) or −80 mV (*Trichoplax* AKDF^19383^) with an Oocyte Clamp OC-725D amplifier (Warner Instruments), Axon Digidata 1550B digitizer (Molecular Devices), and current was sampled at 500 or 1000 Hz and low-pass filtered at 100 or 200 Hz. Current amplitudes were measured in Clampfit 11.1 (Molecular Devices) including additional offline 20 Hz filtering (Bessel 8-pole) for display in figures. Time course of MK-801 block was quantified with first order exponential fits in Clampfit 11.1.

Subsequent data analyses were performed in Prism v9 (GraphPad Software). For concentration-response data, peak current amplitude or peak decrease in current amplitude was plotted against ligand concentration and fitted with four parameter non-linear regression for each recording. These were averaged to give means ± SEM in main text, and a single fit to the average normalized responses (±SEM) is shown in figures.

Schild analysis ^30^ in Fig. 2D and Fig. S2 was performed according to Frizell et al. ^75^. Log current responses to two (or three) concentrations of agonist in the absence of DCKA were fit with a three-parameter Hill equation (Graphpad Prism). This slope was used to constrain similar fits to log current responses to two concentrations of agonist in the presence of increasing DCKA concentrations. These fits were compared to the first control fit (Fig. S2), yielding concentration ratios. (Concentration ratio −1) was then plot against DCKA concentration, and the y = 1 intercept was used to identify pA_2_, which is equivalent to K_B_ when slope = 1. We did not constrain slope here, yielding slopes close to unity for WT and >> 1 for mutants.

### Bioinformatic analyses

In generating a phylogenetic tree covering the broad superfamily of iGluRs in animals, we first assembled iGluR subunit amino acid sequences by blasting with NMDA receptor (rat GluN1) and AMPA receptor (rat GluA2) against two ctenophores (*Euplokamis dunlapae*, *Mnemiopsis leidyi*), one poriferan (*Oscarella carmela*), two cnidarians (*Aurelia aurita* - Roscoff strain, *Nematostella vectensis*), one placozoan (*Trichoplax adhaerens* – Grell strain), and, from bilaterians, two xenacoelomorphs (*Hofstenia miamia*, *Xenoturbella bocki*), two protostomes (*Crassostrea gigas*, *Lingula anatina*), three invertebrate dueterostomes (*Acanthaster planci*, *Saccoglossus kowalevskii*, *Branchiostoma belcheri*) and one vertebrate deuterostome (*Rattus norvegicus*) using published xenacoelomorph transcriptomes ^76^, the OIST Marine Genomics Unit database (*Aurelia aurita*, *Lingula anatina*), Joint Genome Institute (*Nematostella vectensis*), ENSEMBL Metazoa (*Mnemiopsis leidyi*) and NCBI (all others, except rat, from UniProt). Retrieved amino acid sequences lacking ligand-binding domain or channel-forming helices were excluded, duplicates of over 85% identity were excluded, and 212 remaining sequences were aligned in MAFFT ^77^ along with *Arabidopsis thaliana* (plant) GLR 3.4 as an outgroup ^78^, and this alignment was used to generate a GLR 3.4-rooted phylogeny with FastTree ^79^. The alignment and phylogeny can be freely downloaded from https://www.lynaghlab.com/resources. MAFFT and FastTree were used as plugins within Geneious Prime 2023 (Dotmatics). Secondary structure and relative position of amino acids in alignment extraction (Fig. 5C) are based on published NMDA receptor and AMPA receptor X-ray structures^25,80^.

The analysis of placozoan genes in Fig. S4 and Supplemental text utilized the *Trichoplax adhaerens* (“Grell”) genome ^40^ and the *Trichoplax* sp. H2 (“Panama”) genome ^39^, both retrieved from NCBI, and the *Hoilungia hongkongensis* genome ^38^ retrieved from https://bitbucket.org/molpalmuc/hoilungia-genome/src/master/.

## ACKNOWLEDGEMENTS

This project has received funding from the European Research Council (ERC) under the European Union’s Horizon 2020 research and innovation programme (grant agreement no. 803714) and from The Research Council of Norway (project no. 234817). The authors are grateful to Adriana Alvizu (Michael Sars Centre) for help with molecular biology and David MacLean (University of Rochester Medical Center) for plasmids.

## AUTHOR CONTRIBUTIONS

Conceptualization, SS, TL; Methodology, SS, OS, YW, HYH, TL; Investigation, SS, OS, YW, HYH; Resources, HYH; Writing – original draft, OS, HYH, TL; Writing – review & editing, SS, OS, YW, HYH, TL; Visualization, OS, TL; Funding acquisition, TL.

