## Supplementary Figures and Text for "Constitutive activity of ionotropic glutamate receptors via hydrophobic substitutions in the ligand-binding domain"

**CONTENTS**

| **Figure S1.** Expression of NMDA receptors incorporating α-hydroxy amino acids via nonsense suppression. | **Page 2** |
| --- | --- |
| **Figure S2.** Schild analysis of ligand-current inhibition by DCKA. | **Page 3** |
| **Figure S3.** GluN1(4a)-containing receptors are similar to GluN1(1a)-containing receptors. | **Page 4** |
| **Figure S4.** A placozoan AKDF iGluR gene with a tyrosine residue at the αI D732 position. | **Page 5** |
| **Figure S5.** Ditetrabutylammonium Nvoc-Val-OpdCpA | **Page 6** |
| **Figure S6.** Ditetrabutylammonium Vah-OpdCpA | **Page 7** |
| **Figure S7.** Ditetrabutylammonium Nvoc-Ala-OpdCpA | **Page 8** |
| **Figure S8.** Ditetrabutylammonium Aah-OpdCpA | **Page 9** |
| **Supplemental text** | **Page 10** |
| **Supplemental information references** | **Page 11** |

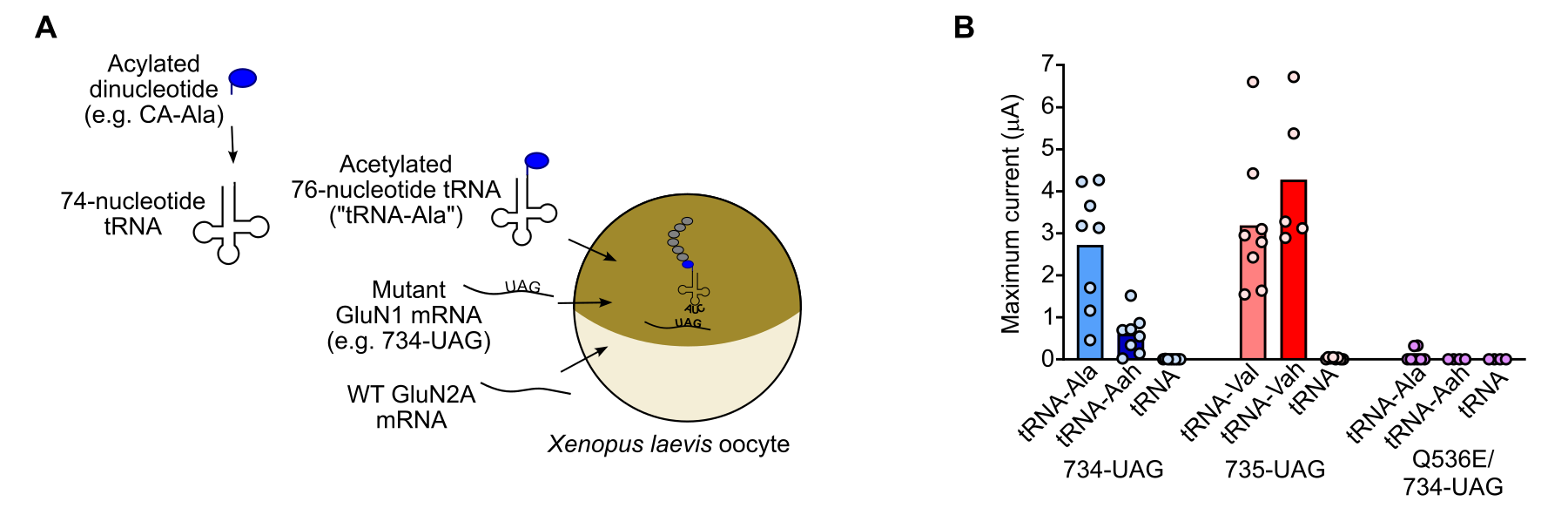

**Figure S1. Expression of NMDA receptors incorporating α-hydroxy amino acids via nonsense suppression.**

(**A**) Acetylated dinucleotide was enzymatically ligated to THG73 tRNA lacking the final two nucleotides (1). The resulting acetylated tRNA was co-injected into oocytes along with UAG mutant GluN1 and WT GluN2A mRNAs. This was performed for alanine (Ala), valine (Val), and α-hydroxy acids lactic acid (“Ala”) and 2-hydroxy, 3-methyl butyric acid (“Vah”), as well as the “empty” dinucleotide, as a control for acylation of spent tRNAs with endogenous amino acids.

(**B**) Summary data (column, mean; dots, individual data points; n = 5-8) of glycine (100 µM) + glutamate (100 µM) gated currents in *Xenopus laevis* oocytes after injection of WT GluN2A mRNA, mutant GluN1 mRNA, and either regular amino acid-tRNA, α-hydroxy acid analogue-tRNA, or empty tRNA as a control for non-specific incorporation of endogenous amino acids.

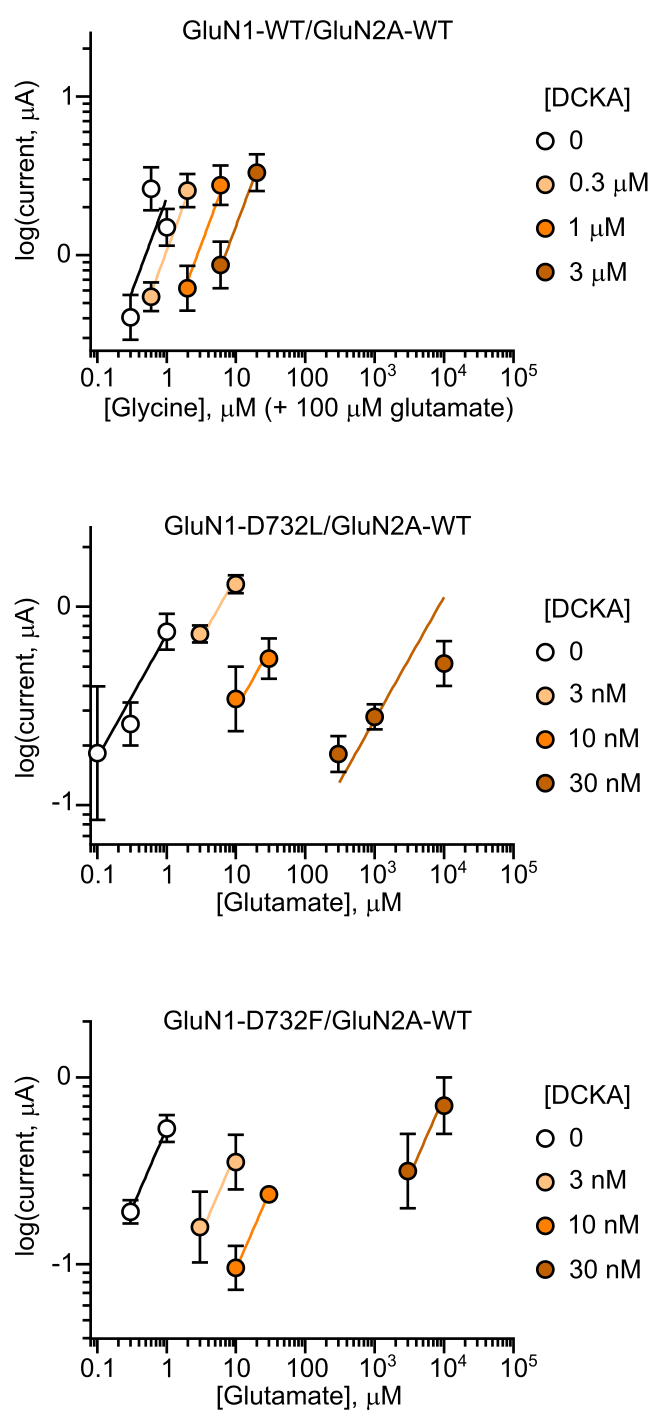

**Figure S2. Schild analysis of ligand-current inhibition by DCKA.**

Log current responses to two (or three) concentrations of agonist in the absence of DCKA were fit with a three-parameter Hill equation (Graphpad Prism) yielding a slope that was used to constrain subsequent fits to log current responses to two concentrations of agonist in the presence of indicated DCKA concentrations. From these slopes, concentration ratios were established and plot against DCKA concentration in Figure 2D.

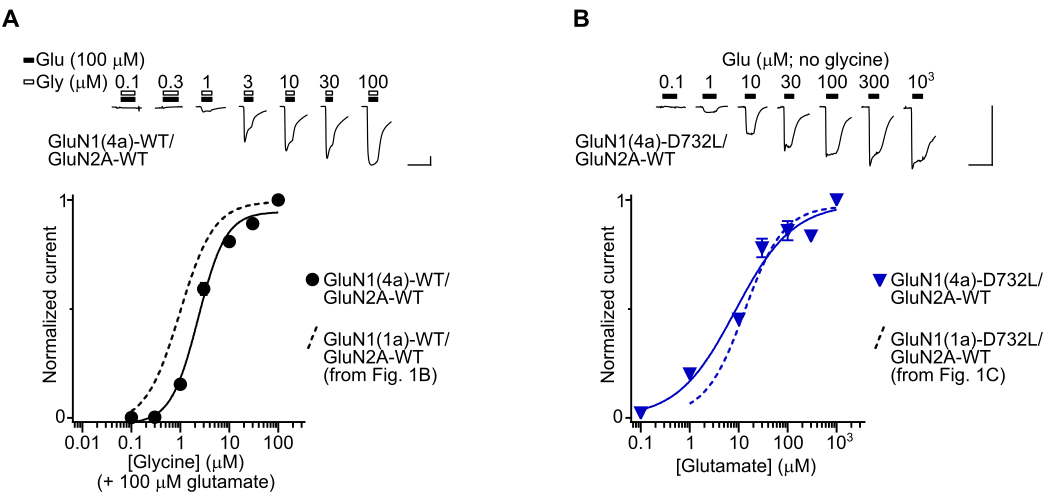

**Figure S3. GluN1(4a)-containing receptors are similar to GluN1(1a)-containing receptors.**

(**A**) *Upper*, example current responses to increasing glycine concentrations in the presence of 100 µM glutamate in an oocyte expressing WT GluN1(4a)/GluN2A receptors. *Lower*, mean ± SEM (n = 6) normalized current responses to increasing glycine concentrations in WT GluN1(4a)/GluN2A receptors.

(**B**) *Upper*, example current responses to glutamate in the absence of glycine in an oocyte expressing GluN1(4a)-D732L/GluN2A-WT receptors. *Lower*, mean ± SEM (n = 6) normalized current responses to increasing glutamate concentrations in GluN1(4a)-D732L/GluN2A-WT receptors.

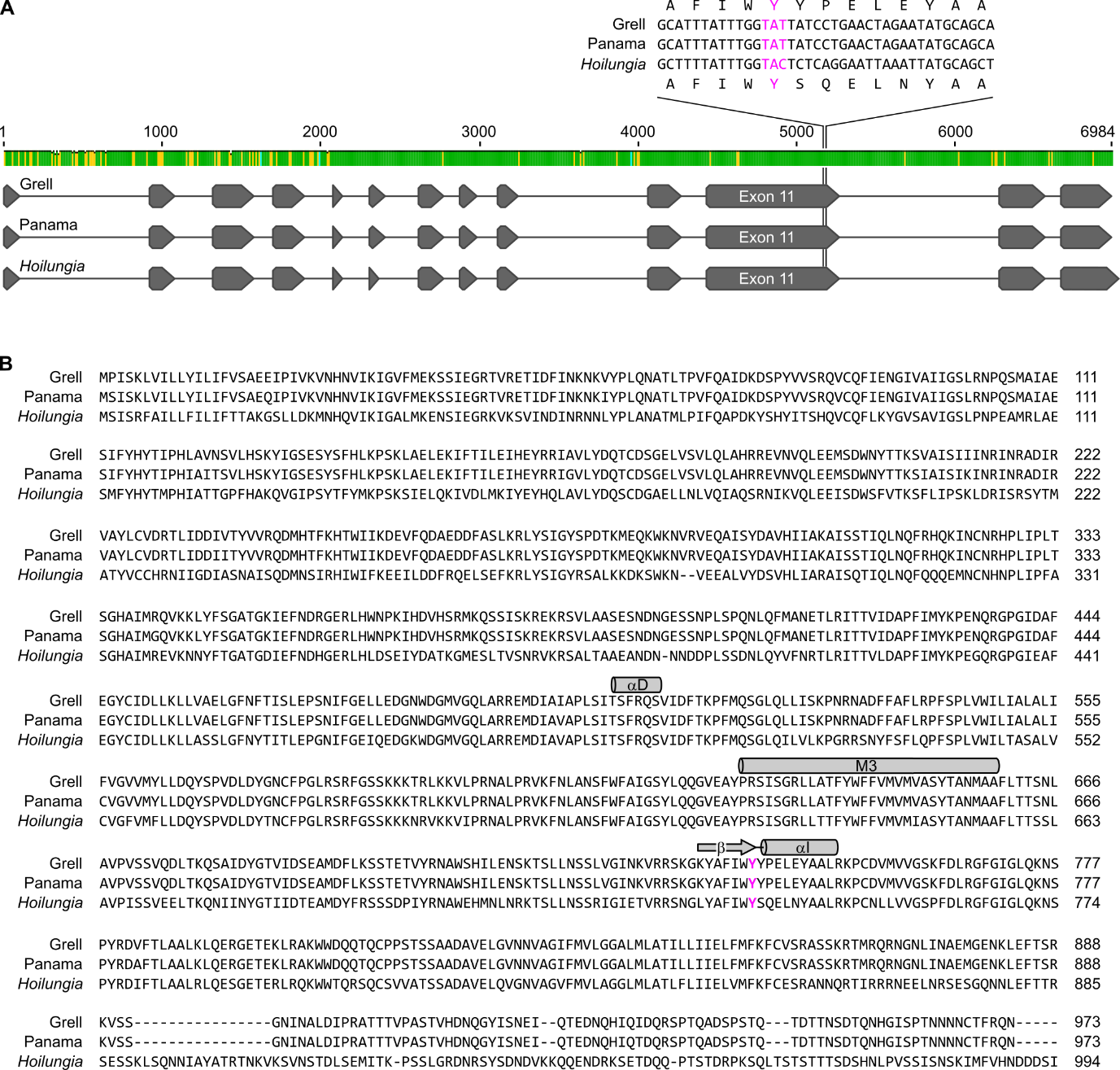

**Figure S4. A placozoan AKDF iGluR gene with a tyrosine residue at the D732 position.**

(**A**) Alignment of our gene annotations (exons grey boxes, introns grey lines) of *Trichoplax adhaerens* (“Grell”) gene TRIADDRAFT_19383/scaffold NW_002060943.1, *Trichoplax* sp. H2 (“Panama”) scaffold NOWV01000306.1, and *Hoilungia hongkongensis* Contig 12. Green nucleotide alignment score above (green, identical; yellow, single nucleotide difference; cyan, indel) refers to alignment of only Grell and Panama genes. Magnified view of Y742 (pink) region shown above. Grell gene corresponds to Genbank entry XM_002107744, an incomplete mRNA transcript; we annotated the complete gene in scaffold NW_002060943.1 based on RDD36848.1 – an automated annotation of the Panama gene – typical GT/AG splice sites, and alignment with other iGluRs. Our annotation of the Panama gene corresponds to automated annotation RDD36848.1, but we excluded a small exon between exons 9 and 10 because it is not present in the corresponding Grell mRNA transcript of TRIADDRAFT_19383 and it would lead to a large, unusual insertion in the iGluR ligand-binding domain. Our annotation of *Hoilungia* Contig 12 was based on an mRNA transcript (TR3071_c1_g1_i1) from the same publication (2) and alignment with other genes.

(**B**) Amino acid sequence alignment of the above genes. Selected secondary structures described in main text are indicated by cylinders (α-helices) or arrows (β-strands) according to homology with published AMPA iGluR structures (3). Y742 (Grell gene numbering) coloured pink.

| 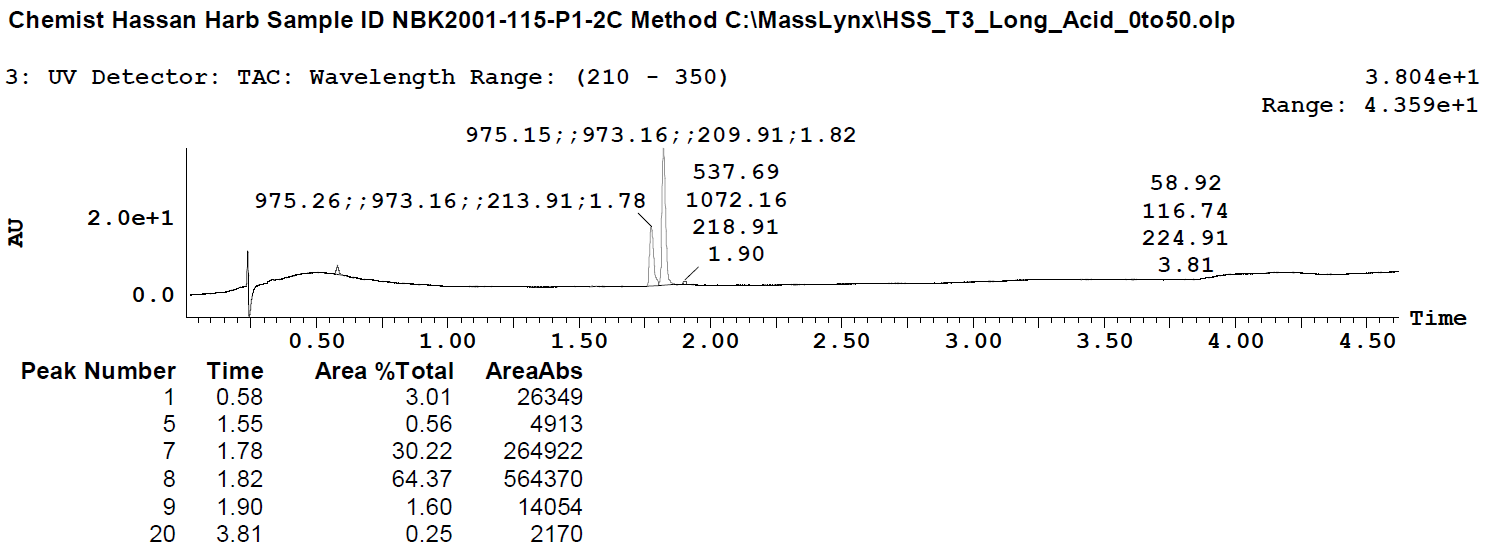 | |
| --- | --- |
| 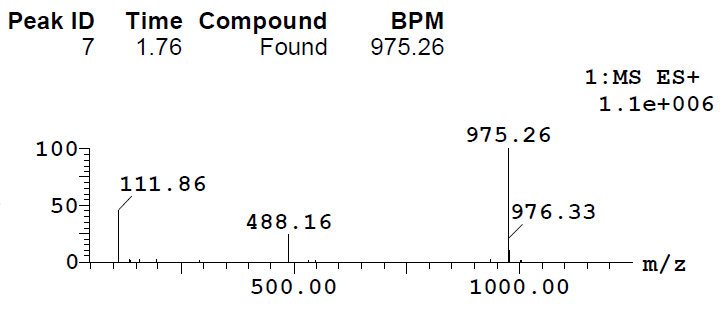 | 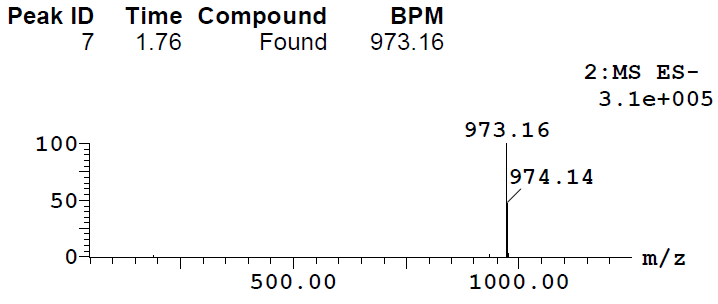 |
| 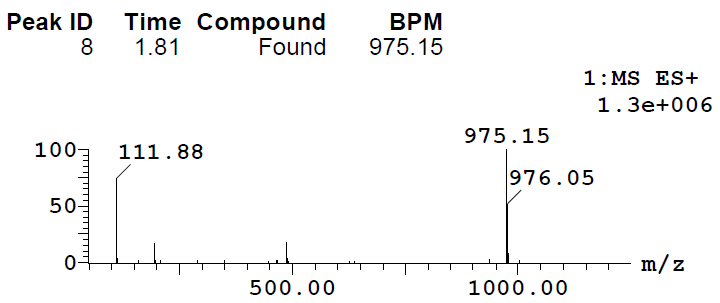 | 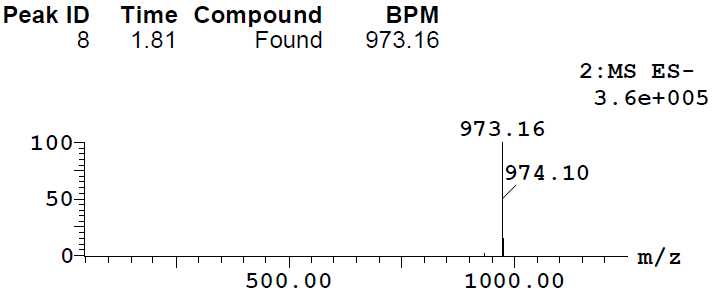 |

**Figure S5.** **Ditetrabutylammonium Nvoc-Val-OpdCpA**

UPLC-MS results.

| 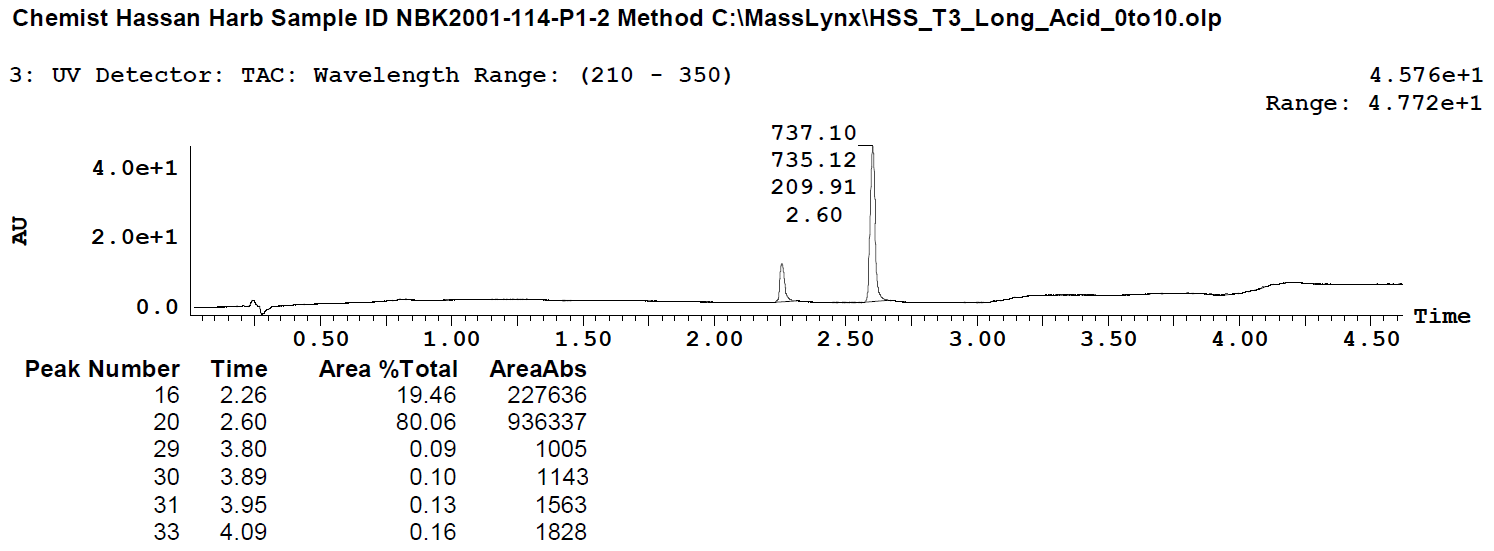 | |
| --- | --- |
| 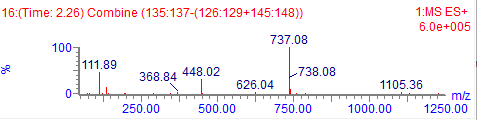 | 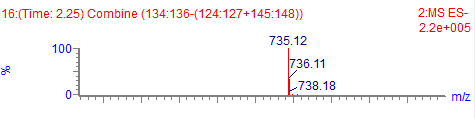 |
| 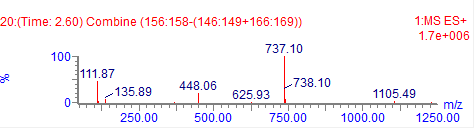 | 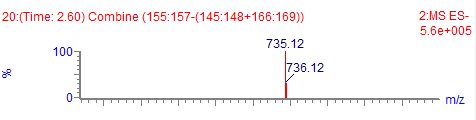 |

**Figure S6. Ditetrabutylammonium Nvoc-Vah-OpdCpA**

UPLC-MS results.

| 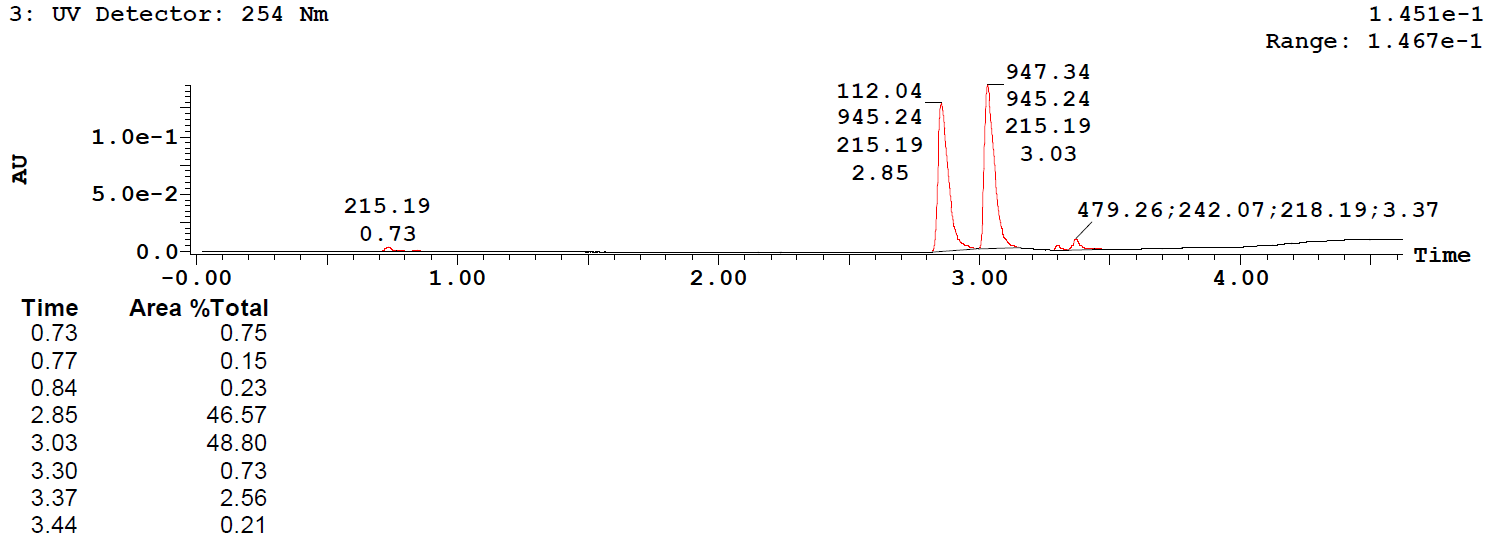 | |
| --- | --- |
| 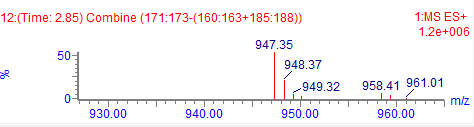 | 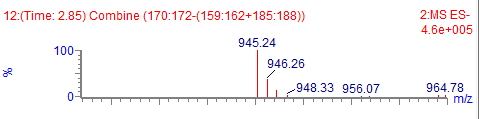 |
| 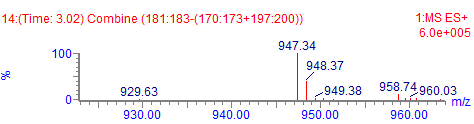 | 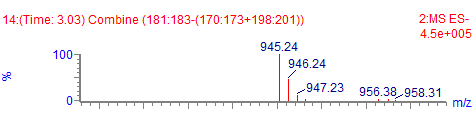 |

**Figure S7. Ditetrabutylammonium Nvoc-Ala-OpdCpA**

UPLC-MS results.

| 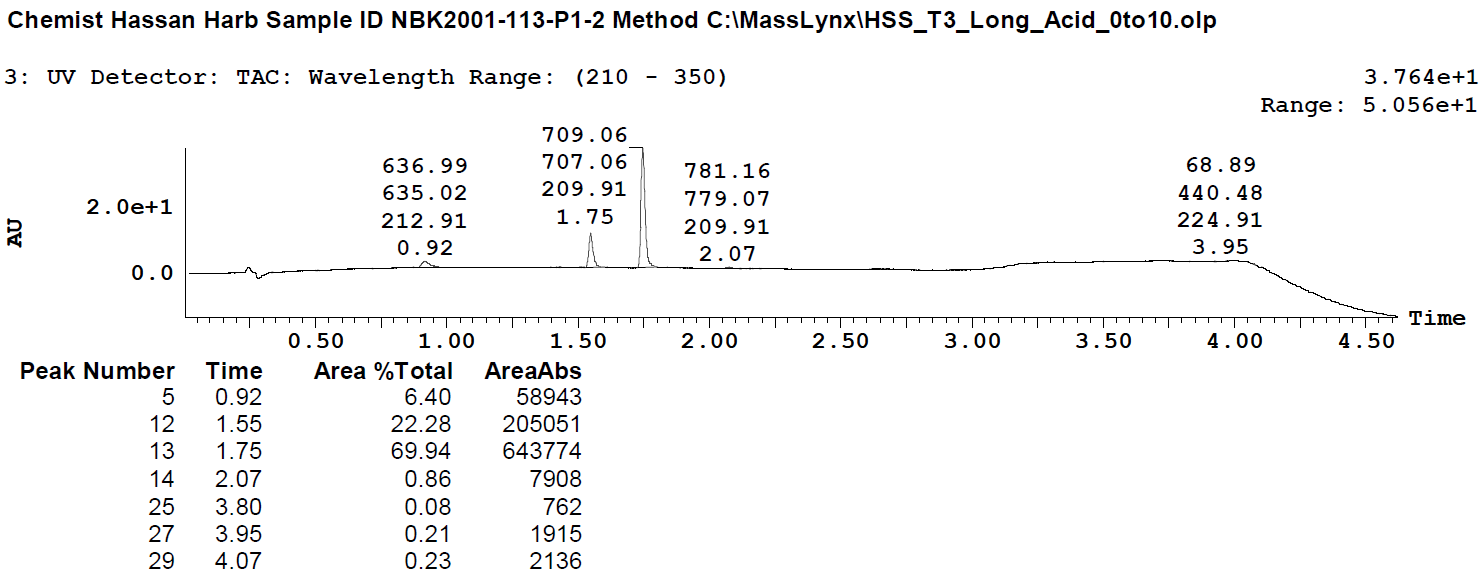 | |
| --- | --- |
|  | 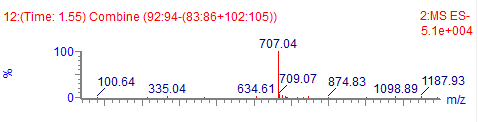 |
| 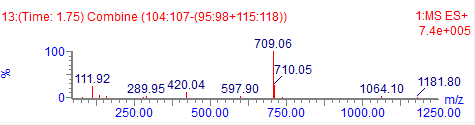 | 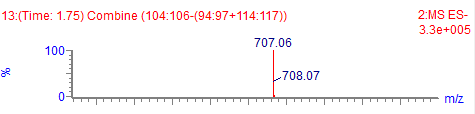 |

**Figure S8. Ditetrabutylammonium Aah-OpdCpA**

UPLC-MS results.

**Supplemental text**

Annotation of AKDF^19383^ in *Trichoplax adhaerens* (“Grell”) scaffold NW_002060943.1

*Black*, exons. *Blue*, introns.

ATGCCAATATCCAAATTGGTTATCTTATTATATATTCTTATATTTGTTTCCGCTGAGGAAATCCCAATCGTAAAAGTAAATCATAACGTCATTAAAATTGGTAAGCCTAATAAAATAGGAAGTTTATGACATGAATGATATTATAGTTACTTATTGGTGCTAGGTGTCTACGTTAATTCTTCGATAACTAATAACCAACTTCCTAACCAACAGCTAATCAGAGTAAATTTAATTACGAGTGAAATGGTAACTTATTTTAAATCAAGGCAGTATTTTTAGCATAACGGCGATATTATACTTATTGTTTATGATCGATAACTTGATTATTGTTTAATATTTTCCATTAGTGAGCTGCACTAATAAGATTTCAAAATTAAAGCTAGTTAGATCCTGCTATCATCGTTCAATACTTTTCTGCAGTGGACTGAATTAATAAGATTTCAGTATTAAATTATAGCTACTCGAAATCTGAATACTGATATTTACTTCTTGGAAGCGATAGTAAAGAATGAGTAACTTTTATAGTAAAACCACTAAAAGTTAAGTTGCGTGATTGTTCGCTAAACATTTACTGTAATAGTAGAGGATCTACCATTTGACTTATGGTATAATTTTATTACGCGGAGTATTCGATTTAATATGATACCTGGTGCCAGTTACTAAAAATTTTTTTCATTCGATCTATAGTATCTACTGACTAACTCATAACAAAGAAATTACGTAGCTGTAACTGTTAAATTTTATCTTAACTATGCTGATTATTGACGTATTATTGAATTAATCGGTGGCAATAATGTCTATAATTATAATTTTAAGCTCAAATAGTGCAACTATTAGTTTCATTAGCTTACGTAAGATGTGAAGAAGAATTCGAATTGCTGGATGCAAAGTCGAATTATAATTTTTTAATCATCTTTAGGCGTATTTATGGAAAAATCATCAATTGAGGGACGAACCGTTAGAGAAACCATTGATTTTATCAACAAAAATAAAGTTTATCCATTACAGAATGCTACACTAACGCCAGTCTTTCAGGCTATCGACAAAGACAGCCCTTACGTAGTATCTCGGCAAGGTTAGTGCAGCCTCTAAGATGGAATTTTATTTTATTTAACTATAGTATAACTTTGCACAATAAATCTTGAACAGTAAGTGCAATGTATTTTCGTATAACTCAGATGAAATTATATTAAATGTCAGTATTTTGGATACTTTCTAACTGCATCTAGAACGTGAAATAGATTTGTACTAAAAAATTCTGAATTACCCTTTTTTTCTACTGTTATTAAGTTTTTATGATCGATTATTTTAGTGTGCCAATTTATAGAAAATGGTATTGTAGCTATTATTGGATCGTTAAGAAATCCTCAATCAATGGCTATAGCGGAATCTATTTTTTATCATTATACAATTCCCCATTTAGCTGTCAATTCAGTATTGCATTCTAAATATATTGGTAGCGAATCTTATAGTTTCCACCTGAAGCCATCAAAGTTAGCGGAGTTGGAAAAAATCTTTACCATTTTAGAAATTCACGAATATCGTCGGATTGCAGTTCTATACGATCAAACGTGTGGTTAGTTTGTCGTCCAGTTTATTATTACGATAAGCAAAAATTTAAGGGATTAAGCTACTTGCAGTCAATGTAGTTATGCACCATGAAACTTGTTAAGCTTAACTAATTTTTCCTTTTAGATAGCGGAGAGTTGGTGAGTGTGTTACAGTTAGCACACCGCAGAGAAGTTAATGTTCAACTAGAAGAAATGTCTGATTGGAATTATACGACAAAATCCGTTGCTATCTCAATTATAATTAATCGCATTAATCGCGCTGATATTAGAGTTGCTTATTTATGTGTAGATCGTACCCTCATTGATGATATTGTTACATACGTAAGTAATTTTAATATTTAAGTAAAATAATCAGTTTTTTTTGTTATTTAGGTATACCTAAGGGTAGATTTAATTATGTCACATACGTGTATTCGTTCAATGAAATCACCATTCATTACATGATTTTACGTGAGAAATAGTAATGCGTAAAATTACTGGTTTAACTCATTATTAAACATGCAACAGGTTGTACGGCAAGACATGCATACTTTCAAACATACTTGGATTATTAAAGACGAGGTTAATTCCTATCAATTGTTTATACAATTAAGATTATTTCACTATAAATTGTTCTCTAATCTAAAAACTAACTTTGCCCATCGCATAAGCTTAGCAACTAATCTCTTTATTAGTTAATGTAGCGTTTGGAATGCATTTAATTGTTAATACTGTATTACGAATAACTTCAGGTATTTCAGGATGCAGAAGACGATTTTGCTAGCTTAAAGAGGTTATATAGTATAGGTTACAGCCCAGATACCAAGATGGAACAGAAGTGGAAAAATGTTAGAGTAATGAAAGAATTTTCGTTTATGTTTTTATTTTATTTAATGTAATTATTTCCTGATTATAGTACTCGTATAGGCATATATTGCAATTTAAATTCCTAGTGGTTTTAAATTAAAAAGCGAATTACAAGTGTATCTTTACATAATATACTGCTGATACTTTATGATTTAATGCACAATGGTATTCTCTTATTTCGGATTTCCTTCCACTTAGGTTGAACAAGCTATCTCGTATGATGCGGTCCACATTATTGCAAAAGCTATCAGTTCGACTATCCAATTAAATCAATTTCGTCATCAGAAAATTAATTGCAATCGTCATCCACTAATTCCATTGACATCCGGTCATGCTATTATGAGGCAGGTTAAAAAGGTGAATATAAAATGAGTCAATGGGCACACAGCAGCTTGTAAGAAGTTTGCTTTAAAAGCTTATTAATTATTAATTCAAGCTGATTATTGACTATCACAGCTTTATTTCTCTGGAGCTACGGGTAAAATTGAATTTAATGACCGCGGAGAAAGATTGCATTGGAATCCTAAAATACATGATGTTCATAGTCGAATGAAGCAATCTTCGGTAATTTCAAATTAAATCCTTGTATATTCAATTAAACGATATAGTACATTCAAAGATTCACACCAGTTGTCGAAATACGTAGTAACAGTGATCTCGCCTGTTATTTACGATCTTGAACGCTTCATACAGATTTCAAAGCGTGAGAAGCGAAGTGTACTTGCGGCTTCCGAAAGTAATGATAATGGTGAATCTTCGAATCCTTTAAGTCCGCAAAATTTACAGTTTATGGCCAATGAAACTTTGCGAATTACCACAGTGATAGTGAGTATAACGTACGATCTGCTATCATGCCGTGATTGATATCGATAACGGGTTTTATTTACTTTGCTAATGAATTCAAATTTTAAATTCGATAATTAATTTAATAGGATATAACTACCTCCTCAGATATTAGTTTGAATATGAGTGAAACTTTTTGTAAGCATTTGGCACAAATTCATAATTACATTATTTGTCCTCTAGTTCGCGTTACTATTTTTAAGCTATGAACGCACCTCATATTGGTAGGCAGTTGTTTAAAATATGATGCACTACTCGTATTATGGCTGGAGTTGTTTCTTAAACTTGTTAGTATTACTTTACAGTATACTGAATGTTATTATTTGCCTTGCTAGTGAACTCAAATTTGGATTGTGATAATTAATTTTAATAAGATGTAATTACCTTTCAGATGTCAGTTCAATATAAGTGAAATCTGTTTCGTAAGCATTTGACATAAATTATAATTATAGTGTGCTCTAGCCCATGATACTAGTTTTGATCCATGAAGAGCATATATCGTAAGGCACCACATATTTATTTTTAGGCCCATTGATTTTAAAACATGGTACGGTACTGTTTCATTACCAGTGTTGTCTATTTGATCTGGTGTTATTGCATAAGGGACAACAGAATCTATTAATTTTGTTGAGCATGAATAATTTTACCAACATATAAGAGAAAATCTATCGTACATACTCTAAAAATCAATATTTACTATTTCAGTAATCAAATAGAGTTTCAATTCATTCTGATCATGAGTATATTTCTGTTTTAACTGTTTCCTTTCTCCTTAAGTACTTATTTGTGAATATCGTTTAGGATGCACCGTTTATTATGTACAAACCAGAAAATCAAAGAGGGCCTGGAATCGATGCATTCGAAGGCTATTGCATTGACTTATTAAAATTATTAGTTGCAGAACTAGGCTTCAACTTTACCATTAGTTTAGAACCTAGTAATATCTTTGGAGAGTTATTAGAAGACGGCAACTGGGACGGTATGGTTGGCCAGTTAGCACGAAGGGTACATAAATTTAAAATCAGTGATCGCTTCCATTTAATTTTTCTCTCTTAATGATTATCTATCATATCTTAGAATATTACACCAAGGACCTAGTAAATTTGATTGCATTAACAAATTGTAATTCCTATACTAAAATAACTATTCTTGTATTTCATTTGAATAGGAAATGGATATAGCAATAGCTCCATTATCTATTACGTCGTTTAGACAAAGTGTAATTGATTTTACTAAACCGTTTATGCAGTCAGGATTGCAATTATTAATTTCGAAGCCCAATCGAAATGCGGATTTTTTCGCTTTTTTAAGACCATTCTCACCGCTTGTCTGGATTTTGATTGCTTTAGCTTTAATTTTTGTCGGCGTAGTGATGTATTTATTAGATCAATATAGTCCGGTCGATCTTGATTATGGTAATTGCTTCCCTGGCTTGAGATCTCGTTTTGGTAGTAGTAAAAAGAAAACTCGACTTAAGAAAGTTCTACCTCGTAATGCTTTACCCCGTGTTAAATTTAACTTAGCGAACAGTTTCTGGTTTGCAATTGGTTCTTATTTACAGCAAGGAGTTGAAGCTTATCCTCGATCTATTTCTGGAAGATTATTAGCTACATTTTATTGGTTTTTCGTTATGGTTATGGTTGCATCTTATACAGCGAATATGGCTGCTTTTTTGACCACTTCTAATTTAGCTGTTCCTGTATCTTCAGTCCAAGATTTAACTAAGCAAAGCGCCATCGATTATGGAACAGTGATTGATAGCGAAGCCATGGATTTTTTAAAATCATCGACCGAAACAGTCTACAGAAACGCTTGGTCTCATATTTTAGAAAATAGCAAGACTTCTTTACTTAATTCATCACTTGTTGGTATAAACAAAGTCAGGCGAAGTAAAGGGAAATATGCATTTATTTGGTATTATCCTGAACTAGAATATGCAGCATTGCGAAAACCTTGCGATGTTATGGTAGTTGGTTCAAAATTTGATCTACGCGGCTTTGGTATTGGTAAGTCTGATATCAATAGCCAAAAGTACGAACGATTGCTTTAATTGTTAGATAATCGTGATACAAATTATCCATCTTTATTTAGCTAAGAAGTATACTGATACTATATAGTATTGAATTAGTACTTAAGGTTTTATATTATTATGCTATTCACAGGGTGCAGTAAAGAATTTATTATTACAATAGTTACTCTAATATTTCCTATTTTAGCAATTGTAAGGTATTAGTAATTGACTAGATTATGAACGTGAAAAATTCCTGTCTAGCACTTCCTTGGTTAAGGAAATATTGCTTGAATGAGATTGCTACTATTAGCACTTAAAAAGTCGTATACGGTTATGTAATTTAGCTAAAAAAATAGTGCTAAATTATAAAAGAATTTTTCGATATTGATGACTTTAACATTTCCTGCTTTAGCAGTTTCAAATGTATCAATAATTGACCTGGTAGTTTATCTGATAGTACCCTTTCTAGCAAATACTATATAGTCCCGTATCTGTTGCTTACGTATCGTATTTGTTAGGCATTGTATATGACAAAGTATGCAATAACTTTCTAAATATTATGCCATAAATGAAGTTTACGAAATTTTAGTAGTACTAAAGACGGTGTTCTCTACTCTGTCGATAAAATTAATAATCGTTAACTGAATTGTCGATGAGGTAATTTCTGCTTACAAGTTTCACACATATGTCGAGATTGTCGACATGTAGGTAGTTAGCAAAACGCTGCTATTCCAAGTCATGAATATTTATTAACTAGTAAATCAGCACGTAACATACTTTATGATGATAATTAATTATTTAAATTGCTAGATCATGTATGAAGTTGCAAACTCCTAGACATTATTACTCTAGCGATTTTTTCTAATTAATAATTAAGAGCAATAGCTTAGGAAGTTGATGTGATAAATTTTTGTATAGCTAGTTCATAATTGTATAGATAAGTGTACTAATTGCTATGGCAGATTCAAAGCACTTTAAGATTGCGTACTTACAATCATCAGGTTTACAGAAAAACTCTCCATATAGAGATGTCTTTACCTTAGCTGCCTTAAAATTGCAAGAAAGAGGAGAGACAGAAAAGCTGCGAGCAAAATGGTGGGATCAGCAGACTCAGTGCCCTCCTTCAACGTCTTCTGCAGCTGATGCAGTTGAATTGGGGGTCAACAATGTAGCTGGTATCTTCATGGTTCTTGGTGGTGCTTTGATGTTGGCTACCATCCTTCTGATCATTGAACTCTTCATGTTTAAATTTTGCGTATCTCGTGCCAGTAGCAAACGAACCATGAGACAGGTTTGTTTAGATAGATACTTCAAATAAATTAGCACTTCGTTTCGAATATCATATGACCATCAAGGTGATTTCCCATATTGATCTACAATTTTCTATTACAGAGAAATGGAAACTTGATTAATGCAGAAATGGGTGAAAATAAATTAGAATTTACTTCTAGAAAAGTCAGCTCTGGTAATATAAATGCTCTTGATATACCCAGAGCGACGACAACAGTTCCTGCAAGTACAGTTCATGATAACCAGGGTTACATAAGCAATGAAATACAAACAGAAGACAACCAACATATACAAATCGACCAGCGATCACCTACTCAAGCTGACTCACCATCGACCCAGACAGACACGACTAATTCAGACACACAAAACCACGGTATTAGTCCTACTAATAATAACAATTGCACTTTTAGGCAAAACTAA

Annotation of AKDF^19383^ orthologue in *Trichoplax* sp. H2 (“Panama”) scaffold NOWV01000306.1

*Black*, exons. *Blue*, introns. *Magenta*, potential additional exon.

ATGTCAATATCCAAATTGGTTATCTTATTATATATTCTTATATTTGTTTCCGCTGAGCAAATCCCAATCGTAAAAGTAAATCATAACGTCATTAAAATCGGTAAGCCTAATAAAATAGGAAGTTTATGACATAAATGATATTATAGTTACTTATTGGTGCTAAGTGTCTACCTTAATTCTTCGATAACTAATAACCAACTTCCTAACCAACAGCTAATGAGAGTAAATTTAATTACGAGTGAAATGGTAACTTATTTTAAATCAAGGCAGTATTTTTAGCATAACGGCGATATTATACTTGTCGTTTATGATCGATAACGTGACTATCGTTTAATATTTTCCCGCAGTGAGCTACACTAATAAGATTTCAAAATTAAAGCTAGTTAGATCCTGCTATCATCGTTCAATACTTTTCTGCAGTGGACTGAACTAATAAAATTTCAGTATTAAATTAAAGCCACTTGAAATGTGAATACTGATATTTACTTCTTGGAAGCGATAGTAAAGAATGAGCAAATTCTATAGTAAAACCACTAAAAGTTAGGTTGTGTGATTGTTCGCTTAACATTTACTACAATAGTAGAGGATCTACCATTTGACTTATGGTACAATTTTATTACGCGGAGTATTCGATTTAATATCATAGCTGGTGCCAGTTACTAAAAATTTTTTTCATTCGATCTATAGTATCTACTGACTAACTCATAACAAAGAAATTACGTAGCTGTAACTGTTAAATTTTATCTTAACTATGCTGATTATTGACGTATTATTGAATTAATCGGTGGCAATAATGTCTATAATTATAATTTTAAGTTCAAATAATGCAACTATTAGTTTCATTAGCTTACGTAAGATGTGAAGAAGAATTCGAATTGCTGGATGCAAAGTCGAATTATCATTTTTTAATCATCTTTAGGCGTATTTATGGAAAAATCATCAATTGAGGGACGAACCGTTAGAGAAACCATTGACTTCATCAACAAAAATAAAATTTATCCGTTACAGAATGCTACACTAACGCCAGTCTTTCAGGCTATCGACAAAGACAGCCCTTACGTAGTATCTCGGCAAGGTTAGTGCAGCCTCTAAGATGGAATTTTATTTTATTTAACTATAGTATAACTTTGCACAATAAATCTTGAACTGTAAGTGCAATGTATTTTCGTATAATTCAGATGTAATTATTATTAAATGTCAGTATTTTGGATACTTTCTAACTGCACCTAGAACGTGAAATAGATTTGTACTAAAAAATTCTGAATTACCCTTTTTTTCTACTGTTATTAAGTTTTTATGATCGATTATTTTAGTGTGCCAATTTATAGAAAATGGAATTGTAGCTATTATAGGATCATTAAGAAATCCTCAATCAATGGCTATAGCGGAATCTATTTTTTATCATTATACAATTCCCCATATAGCTATCACTTCGGTATTGCATTCTAAATATATTGGTAGCGAATCTTATAGTTTCCACCTGAAGCCATCAAAGTTAGCAGAGTTGGAAAAAATCTTTACCATTTTAGAAATTCACGAATATCGTCGGATTGGAGTTCTATACGATCAAACTTGTGGTTAGTTTGTCGTCCAATTTATTATTACGATAAGCAAAAAATTTAAGGGATTAAGCTACTTGCAGTCAATGTAGTTATGCACCATGAAACTTGTTAAGCTTAAATAATTTTTCCCTTTAGATAGCGGAGAGTTGGTGAGTGTGCTACAGTTAGCACACCGCAGAGAAGTTAATGTTCAACTAGAAGAAATGTCTGATTGGAATTATACGACAAAATCCATTGCTATCTCAATTAAAATTAATCGCATTAATCGCGCTGATATTAGAGTTGCTTATTTATGTGTAGATCGTACCCTCATTGATGATATTATTACATACGTAAGTAATTTTAATATTTAAGTAAAATAATCAGTTTTTTCGTTATTTACGTATACCTAAGGGTAGATTTAATTATGTACATACGTGTATTCGTTCAATGAAATCACCATTCATTACATGATTTTACGTGAGAAATAGTTACGCGTAAAATTATTGGTTTAACTCATTATTAAACATGCAACAGGTTGTACGGCAAGACATGCATACTTTCAAACATACTTGGATTATTAAAGACGAGGTTAATTCCTATCAATTGTTTATACAATTAAGATTATTTCACTATAAATTGTTCTCTAATCTAAAAACTAACTTTGCCCATCGCATAAGCTTAGCAACTAATCTCTTTATTAGTTAATGTAGCGTTTGGAATGCATTTAATTGTTAATACTGTATTACGAATAACTTCAGGTATTTCAGGATGCAGAAGACGATTTTGCTAGCTTAAAGAGGTTATATAGTATAGGTTACAGCCCAGATACCAAGATGGAACAGAAGTGGAAAAATGTTAGAGTAATGAAAGAATTTTCGTTTATGTTTTTATTTTATTTAATGTAATTATTTCCTGATTATAGTACTCGTATAGGCATATATTGCAATTTAAATTCCTAGTGGTTTTAAATTAAAAAGCGAATTACAAGTGTATCTTTACATAATATACTGCTGATACTTTATGATTTAATGCACAATGGTATTCTCTTATTTCGGATTTCCTTCCACTTAGGTTGAACAAGCTATCTCGTATGATGCGGTCCACATTATTGCAAAAGCTATCAGTTCGACTATCCAATTAAATCAATTTCGTCATCAGAAAATTAATTGCAATCGTCATCCACTAATTCCATTGACATCCGGTCATGCTATTATGGGGCAGGTTAAAAAGGTGAATATAAAATGAGTCAATGGGCACACAGCAGCTTGTAAGAAGTTTGCTTTAAAAGCTTATTAATTATTAATTCAAGCTGATTATTGACTATCACAGCTTTATTTCTCTGGAGCTACGGGTAAAATTGAATTTAATGACCGCGGAGAAAGATTGCATTGGAATCCTAAAATACATGATGTTCATAGTCGAATGAAGCAATCTTCGGTAATTTCAAATTAAATCCTTGTATATTCAATTAAACGATATAGTACATTCAAAGATTCACACCAGTTGTCGAAATACGTAGTAACAGTGATCTCGCCTGTTATTTACGATCTTGAACGCTTCATACAGATTTCAAAGCGTGAGAAGCGAAGTGTACTTGCGGCTTCCGAAAGTAATGATAATGGTGAATCTTCGAATCCTTTAAGTCCGCAAAATTTACAGTTTATGGCCAATGAAACTTTGCGAATTACCACAGTGATAGTAAGTATAACGTACGATCTGCTATCATGCCGTGATTGATATCGATAACGGGTTTTATTTACTTTGCTAATGAATTCAAATTTTAAATTCGATAATTAATTTAATAGGATATAACTACCTCCTCAGATATTAGTTTGAATATGAGTGAAACTTTTTGTAAGCATTTGGCACAAATTCATAATTACATTATTTGTCCTCTAGTTCGCGTTACTATTTTTAAGCTATGAACGCACCTCATATTGGTAGGCAGTTGTTTAAAATATGATGCACTACTCGTATTATGGCTGGAGTTGTTTCTTAAACTTGTTAGTATTACTTTACAGTATACTGAATGTTATTATTTGCCATGCTAGTGAACTCAAATTTGGATTGTGATAATTAATTTCAATAAGTTGTAATTACCTTTCAGATATCAGTTCAATATAAGTGAAATCTGTTTCGTAAGCATTTGACATAAATTATAATTATAGTGTGCTCTAGCCCATGATACTAGTTTTGATCCATGAAGAGCATATATCGTAAGGCACCACATATTTATTTTTAGGCCCATTGATTTTAAAACATGGTACGGTACTGTTTCATTACCAGTGTTGTCTATTTGATCTGGTGTTATTACATAAGGGACAACAGAATCTATTAATTTTGTTGAGCATGAATAATTTTACCAACATATAAGAGAAAATCTATCGTACATACTCTAAAAATCAATATTTTACTATTTCAGTAATCAAGTAGAGTTTCAATTCATTCCGACCATGAGTATATTTCTGTTTTAACTGTTTCCTTTCTCCTTAAGTACTTATTTGTGAATATCGTTTAGGATGCACCGTTTATTATGTACAAACCAGAAAATCAAAGAGGGCCTGGAATCGATGCATTCGAAGGCTATTGCATTGACTTATTAAAATTATTAGTTGCAGAACTAGGCTTCAACTTTACCATTAGTTTAGAACCTAGTAATATCTTTGGAGAGTTATTAGAAGACGGCAACTGGGACGGTATGGTTGGCCAGTTAGCACGAAGGGTACATAAATTTAAAATCAGTGATCGCTTCCATTTAATTTTTCTCTCTTAATGATTATCTATCATATCTTAGAATATTACACCAAGGACCTAGTAAATTTGATTGCATTAACAAATTGTAATTCCTATACTAAAATAACTATTCTTGTATTTCATTTGAATAGGAAATGGATATAGCAGTAGCTCCATTATCTATTACGTCGTTTAGACAAAGTGTAATTGATTTTACTAAACCGTTTATGCAGTCAGGATTGCAATTATTAATTTCGAAGCCCAATCGAAATGCGGATTTTTTCGCTTTTTTAAGACCATTCTCACCGCTTGTCTGGATTTTGATTGCTTTAGCTTTAATTTGTGTCGGCGTGGTGATGTATTTATTAGATCAATATAGTCCGGTCGATCTTGATTATGGTAATTGCTTCCCTGGCTTGAGATCTCGTTTTGGTAGTAGTAAAAAGAAAACTCGACTTAAGAAAGTTCTACCTCGTAATGCTTTACCCCGTGTTAAATTTAACTTAGCGAACAGTTTCTGGTTTGCAATTGGTTCTTATTTACAGCAAGGAGTTGAAGCTTATCCTCGATCTATTTCTGGAAGATTATTAGCTACATTTTATTGGTTTTTCGTTATGGTTATGGTTGCATCTTATACAGCGAATATGGCTGCTTTTTTGACCACTTCTAATTTAGCTGTTCCTGTATCTTCAGTCCAAGATTTAACTAAGCAAAGCGCCATCGATTATGGAACAGTGATTGATAGCGAAGCCATGGATTTTTTAAAATCATCGACCGAAACAGTCTACAGAAACGCTTGGTCTCATATTTTAGAAAATAGCAAGACTTCTTTACTTAATTCATCACTTGTTGGTATAAACAAAGTCAGGCGAAGTAAAGGGAAATATGCATTTATTTGGTATTATCCTGAACTAGAATATGCAGCATTGCGAAAACCTTGCGATGTTATGGTAGTTGGTTCAAAATTTGATCTACGCGGCTTTGGTATTGGTAAGTCTGATATCAATAGCCAAAAGTACGAACGATTGCTTTAATTGTTAGATAATCGTGATACAAATTATCCATCTTTATTTAGCTAAGAAGTATACTGATACTATATAGTATTGAATTAGTACTTAAGGTTTTATATTATTATGCTATTCACAGGGTGCAGTAAAGAATTTATTATTACAATAGTTACTCTAATATTTCCTATTTTAGCAATTGTAAGGTATTAGTAATTGACTAGATTATGAACGTGAAAAATTCCTGTCTAGCACTTCCTTGGTTAAGGAAATATTGCTTGAATGAGATTGCTACTATTAGCACTTAAAAAGTCGTATACGGTTATGTAATTTAGCTAAAAAAATAGTGCTAAATTATAAAAGAATTTTTCGATATTGATGACTTTGACATTTCCTGCTTTAGCAGTTTCAAATGTATCAATAATTGACCTGGTAGTTTATCTGATAGTACCCTTTCTAGCAAATACTATATAGTCCCGTATCTGTTGCTTACGTATCGTATTTGTTAGGCATTGTATATGACAAAGTATGCAATAACTTTCTAAATATTATGCCATAAATGAAGTTTACGAAATTTTAGTAGTACTAAAGACGGTGTTCTCTACTCTGTCGATAAAATTAATAATCGTTAACTGAATTGTCGATGAGGTAATTTCTGCTTACAAGTTTCACACATATGTCGAGATTGTCGACATGTAGGTAGTTAGCAAAACGCTGCTATTCCAAGTAATGAATATTTATTAACTAGTAAATCAGCACGTAACATACTTTATGATGATAATTAATTATTTAAATTGCTAGATCATGTATGAAGTTGCAAACTCCTAGACATTATTACTCTAGCGATTTTTTCTAATTAATAATTAAGAGCAATAGCTTAGGAAGTTGATGTGATAAATTTTTGTATAGCTAGTTCATAATTGTATAGATAAGTATACTAATTGCTATGGCAGATTCAAAACACTTTAAGCTTGCGTACTTACAATCATCAGGTTTACAGAAAAACTCTCCATATAGAGATGCCTTTACCTTAGCTGCCTTAAAATTGCAAGAAAGAGGAGAGACAGAAAAGCTGCGAGCAAAATGGTGGGATCAGCAGACTCAGTGCCCTCCTTCAACGTCTTCTGCAGCTGATGCAGTTGAATTGGGGGTCAACAATGTAGCTGGTATCTTCATGGTTCTTGGTGGTGCTTTGATGTTGGCTACCATCCTTCTGATCATTGAACTCTTCATGTTTAAATTTTGCGTATCTCGTGCCAGTAGCAAACGAACCATGAGACAGGTTTGTTTAGATAGATATTTCAAATAAATTAACACTTCGTTTCGAATATCATATGACCATCAAGGTGATTTCCCATATTGATCTACAATTTTCTATTACAGAGAAATGGAAACTTGATTAATGCAGAAATGGGTGAAAATAAATTAGAATTTACTTCTAGAAAAGTCAGCTCTGGTAATATAAATGCTCTTGATATACCCAGAGCGACGACAACAGTTCCTGCAAGTACAGTTCATGATAACCAGGGTTACATAAGCAATGAAATACAAACAGAAGACAACCAACATATACAAACCGACCAGCGATCACCTACTCAAGCTGACTCACCATCGACCCAGACAGACACGACTAATTCAGACACACAAAACCACGGTATTAGTCCTACTAATAATAACAATTGCACTTTTAGGCAAAACTAA

Annotation of AKDF^19383^ orthologue in *Hoilungia hongkongensis* contig 12

*Black*, exons. *Blue*, introns.

ATGTCGATATCAAGATTTGCGATTTTATTATTTATACTCATTTTTACTACGGCCAAAGGATCTTTACTGGATAAAATGAATCATCAAGTCATTAAAATTGGTATGTCATTCAATATAGAATATTATTTTGCTTTCAAATAATTTATAATCATATTAAATGTTAAAGATAAAAAGCGCTAAATCTATCTATTAATACACTGGACGTTAACTCATAATTGAGATACTTTATTGTGATATGGATGTAATTTTATATACTGTAAGTGATTCAATAAAATAAGACAAGAATTTCAACCGACTTGAAAAAATGATACTAGGCATTATTATTTCTTGCTACTTTTATTCATTTAATGTAAATTGATAAAATATACTCATTTAATAAATTACTACTTGAAAGTAATTAAGCATAATATAACTACGTATTTAAATGAGGTCCTTACAAATTTTTAACCAAAAGCTCGTGAAAAGGTGACATTATTAATGATGGATATAAATATCGTAACCAAAACTACTTAAGCTACTACTATAACTTATTTAATCAGTTTAAACTTACTTCTGGTATTCTATACCAGATAAATCGGCAATTAAACAAATACGTATATTAGGGGTAACGTAACGAGTATATATATGAGCATTTTCACATACATTTAATAAAAAAACTTTACAGTTATGATTGTTTTGTATATTTTATTATTAGAGATATTGCATAATGTGTTCTTGCTCTTATCAATGTAAGCATTTTAAAGTTATCAAGTAAGAGAAAATATAACCTTTTATAAAGACATTTTTAAAATATTACGTAGATTATTTCTGCATGGCGCAATCAATTGGCTGGAGATTTCAACCAAGAAACTACATTTATGCTTTACTTATTTAGAAATAGGCTTACTTGTACTATACAGTTCTTGTAATGAAATTCATCATATTATATTGTTTTACACATATTTATAATCTATTTTAATCGCTGTAGTAATACGTTTGAATTTATTCCGTTTTATCTTGCTATGTATATAATCTTATGTGGTATACCATATTAAATTAAAGTTCATTGATACTATGCCGTCATTTCGATAATCATTTCTCTCACCAATATCTATAGTTATAATATTTGAGTTCTACATAAAGAATATTAGCATAAATAATGATATAGACTTAATCGTTATGTAAGCCAACAAACGATGAGTAATCATTTCTTATACATAGCTTGTTTATGCAATTATAATTATTCTCTATTTGCGTTATTTAAATCTTTCTATAGCTCAAATTTCACATAACCTTTATTATTTACTCACCTATATGAATAATACTTCAGTTGTAGATCAATTTAAAATATATCATTAATTTTTATCGAATAAATGATCAGCAAATCCCTAATCGAAATGATACATAGCTTAAATTGATTTTGATTAAACGTAACATTTATCATTGTTAGGTGCATTAATGAAAGAAAATTCAATAGAAGGAAGAAAAGTTAAATCTGTCATCAATGATATTAATAGAAATAATCTTTATCCACTGGCCAATGCAACAATGTTGCCAATTTTTCAAGCTCCTGATAAATATAGTCATTATATTACATCTCACCAGGGTTAGTTTATTGTATTAAATATATTCTTTGTTAAATATGCTGTACTAGTTTAAAGCAATTTTTAATGTTATTACTCTTATATATGATTTAAATTAGACTGTCATCAAATTTGTATGTTAATCAAATTTGCTTGGGTTATTTCAGTTTGTCAATTCTTAAAATATGGAGTTTCAGCTGTTATTGGTTCGCTACCAAATCCTGAAGCAATGAGGCTAGCGGAATCAATGTTTTACCATTACACCATGCCTCATATAGCTACTACAGGACCATTTCATGCCAAGCAAGTTGGTATACCATCTTATACTTTCTACATGAAACCATCTAAATCGATTGAACTTCAAAAAATAGTCGACCTTATGAAAATCTACGAATATCATCAGCTAGCCGTGCTATATGATCAATCCTGTGGTTAGTCTATATATTTTTATTCCATTAATTGACGTATTTGTAAGTAGGAGAATTTAGTTAGCATGTCAGAAATCATCAGATTTAAACACTATAAATTATAAATGTCAGAAATTATCCCATATTGGTAGTTATAGTATCCAACCAGATACTTTTGAATGCTTTCCTCTATATAGTTAGTTAAACACTATGTATTCAAAGTCATTAATTTATAGTTAATAAATTATAATATACACATTTCGTATGTTTAGGTTTCTTTGTTGATATTTTTCAATAATAGTTCAAGTTATATTTAAATAATTTATAATTGACCGTAACATATATAACTGATATTATTCAGATACTATAAATTTTAAACTCTCAAATTATAAATTATAAAAGTATTTATATATTTTCTATAATTAGATATTATGTCAGATATTTTCTTGTGCAATATAAAAACATTCATCTTCACCATTTGTTGTTGTTGTAGATGGTGCAGAATTACTAAATTTAGTTCAGATAGCACAAAGTAGAAATATTAAAGTTCAATTGGAAGAAATCTCGGATTGGAGTTTTGTTACAAAGTCTTTTCTGATCCCATCTAAACTTGATCGTATTAGTCGTTCTTATACGATGGCTACTTACGTATGCTGTCATCGAAATATCATTGGCGATATCGCATCCAATGTGAGTAAATCAATTTATATATTGACGCTACCATGGTTAGATATTCTATATATCCAATTTATATTCTAGGCTATCAGTCAAGATATGAATAGTATTAGACATATCTGGATTTTTAAAGAAGAGGTAAAACTTATATCTTATTATTAACTTTCTACCTGGCATGATTTCAAATGGTCATTGCTACTTTAGTTATTATTAGATTTATGTTTCGAAGCTAGTATCACATCATTTCTGCTTCTAATCTGTGATATATTAGAATTTAATTGTATAATCCTTTTTACTCCCTAGATACTCGACGATTTTCGCCAAGAACTTTCTGAATTTAAGAGGTTGTACAGTATAGGATACAGGTCTGCTTTGAAAAAGGATAAATCTTGGAAAAATGTTAGTCACAAATCTATTTAATTATTTCTTCTTGTAACTGATCGCTTTTGATTGCCCGATGGAAAGAGTATACAGATTATAGATGAACTATTTTTTCAATGTTGTAAAATTAAACTTTGATATGATTACTTTTTTGCCTTTGGCATTTAAAAGTCTTGAAAGATTGATTTACTTATGTACTGTATATTATGAATTCATTTTTGTAATCCAGGTTGAAGAAGCTCTTGTTTATGACAGTGTGCACTTAATTGCACGAGCTATCAGTCAAACTATACAATTAAATCAATTTCAGCAACAGGAAATGAATTGCAATCATAATCCGTTAATACCTTTTGCATCTGGACATGCAATTATGAGAGAGGTTAAAAATGTAAGAATCATTCTATTTCTTGATCAATAAACAATTAGAATACTCGTATAAATCCAGTCATACTTTGTTATAAGCAATAAATGATTAAGCGGTTAGTATGTTACCAAACTGTCTATTTACAGTCACTCATTATATTGTAGAATTATTTCACTGGTGCAACCGGTGATATTGAATTTAATGATCATGGAGAGAGATTACATTTGGATTCAGAGATATATGATGCTACTAAAGGGATGGAAAGTTTGACGGTATCATTTTTAATAATAACTATTACGAATATGTATCTTGGGCTGATTAATTATTACATGTTGTATAGTAAATTATTGTCTACTCTTTCCATGTTTATGTAATAACAGGTTTCTAACCGCGTTAAGCGAAGCGCACTCACAGCAGCTGAAGCTAACGATAATAATAATGATGATCCTCTTTCTTCGGATAATTTACAATATGTGTTTAATAGAACTTTACGCATTACAACTGTATTAGTAAGTATCTTTATATAATGCTACATAAAAATAGTGGTAAACTGTTGAGAAATAATTACTAATCAGTTTAGATATATTTAATGTATACGAAGGTATCATATGTAAATGTTATCATCAATATTACCCTAATTTTCTAAAGATAGATTACCATTTCATATTCTTACCCGTATATATATAGTGATTGCATATTCAGTTACTATGATACTATGTGTATATATTTGATGACACATTAACGACATCAAATTGATGAGGAAAATTAAATAAGTACCTTAGTATTGCCACTAATTAATATTTTAATTAGGTAAACTAATATATTAAATAGATAGAAATATATTATATTGATATTTTAGACTACTATTTAGCCATTTATAAAAATATTTTTGTTTAATTTGTGAAATAATTACAGCTGTCAAGGTAATTGACTGAATGAGTGATTGAGTTTCATAGTTGAAAAACAAATGTATATATTTATTGCATAATTGTAGTAGTCGTAATAATTTATTGGTTATACAATAGATCCAAAGTAATGTCGACAGTTTATACGAGTATGACAGTATCTATACTGATCAATTGTTTAATACATGCTAGGATGCACCGTTTATCATGTATAAGCCTGAAGGGCAAAGAGGGCCTGGAATTGAAGCATTTGAGGGTTATTGTATCGATTTATTAAAGTTATTAGCCAGTAGCCTAGGCTTCAACTATACTATTACGTTGGAACCTGGAAATATATTTGGAGAAATACAAGAAGATGGCAAATGGGATGGAATGGTTGGCCAATTAGCACGAAGGGTATGACTGTACTAGTCATTGAAACATGAGCTGGAAATAACTTCAATATTATCGTTATTCTACATTAAAATTAGATTCAAATAAATAAAATATAGAATTCGGTTAACTTCAATGGTATTTACTGTCGTTATGAGTTTCAAATTGAGATTGCAGTTATCAGAGTACTCTGGTAGTTACCTTAACTTATACGTTCAAAATAATCTTATTCTGATACAGCTTAGTATCTAATATAAATAGCTAACATATGGAAAGAATATACTCTATTTTTTTACTCTTTAGTAACTCGTTCTTGTCTTTGTTAATTCAATCTTCATAGGAAATGGACATTGCTGTTGCACCTTTATCTATTACATCTTTTCGACAAAGTGTAATCGATTTTACAAAGCCGTTTATGCAATCGGGATTGCAAATATTGGTTTTAAAGCCTGGTCGGAGATCGAATTATTTCTCCTTTTTGCAACCGTTTTCGCCACTTGTTTGGATATTAACGGCGAGTGCTTTGGTGTGTGTAGGCTTTGTAATGTTTCTTTTAGATCAATACAGTCCAGTCGATCTCGATTATACCAATTGTTTCCCTGGTTTAAGATCACGCTTTGGTAGTAGTAAAAAGAAAAATCGTGTCAAGAAAGTCATTCCACGCAATGCTTTACCTCGTGTTAAATTCAATCTAGCTAATAGCTTTTGGTTTGCTATTGGATCGTATTTACAGCAAGGTGTCGAAGCATATCCTCGATCTATCTCTGGCCGATTATTAACCACATTTTATTGGTTTTTTGTCATGGTCATGATTGCATCTTATACAGCTAATATGGCAGCATTTTTAACAACGTCTAGTTTAGCTGTTCCAATTTCATCGGTCGAAGAATTAACTAAACAAAATATTATCAATTACGGGACTATTATCGATACTGAAGCTATGGATTATTTTAGATCATCGAGTGATCCCATCTATAGAAACGCTTGGGAACATATGAACCTGAATAGAAAAACTTCCTTACTGAATTCATCGCGTATTGGGATCGAGACTGTGCGCCGTAGTAACGGCCTGTATGCTTTTATTTGGTACTCTCAGGAATTAAATTATGCAGCTTTGCGAAAACCTTGTAATCTTTTAGTCGTCGGTTCACCATTTGATCTGCGAGGATTCGGAATTGGTGAGTTATTATGCCCTATGAGAAGTAATATCCAATGATTTGGATCCAGCATGATAGATGCCTTCGAATAAACCAGCGTATGCTTAGTATTTCCTCATGAATCTGTTAATCACATTATTGTTAGTCTTATATTGATGCATTTTCAGCTTATTAGATCATGTCTTGATGAACCTGTAGTATTACTAATAAACTACTCGTATGAACAATGATAAGATTAGCCATCATTGTATGCCAGGTTATTCATATTCTACTATATTAGTCGTAGTCATAAGTTTTTAAAGTACTGCTACTAATTCAATGGTTTAATGCTAATAAAAGTTATTCAAAAATTCCTATGATACGTCTGATGCATAAAGCTAGCTTATAATTGCTAAATTATATCAGTAGTACAACCATATATAACGTCCAACGTATAATGAAAAACGATATCTAACCATGATTTTAACCATAATGCTTATTAAAGTATATTGTACCATAATAGGTCTTTGTGATAGTTATGATCGTGAATCCTTGTCATTTGTTAACAATGTGTTAATTAGTCATCGTTGAATTGCGTCGTCATTCCAATATTTATATTTCTAATGCTATATTGTCAATGTCCAACGGAGAATATATATTTACGTTTACATTTATTTAAAGCATATTAATGTGCTATTAGCTGCGGTTGCAATTGAAGTTAGTGTTTTATATGTATCTCGCTTAGCTAGCACCTTCTTTCCCACCGGTTATTGCTATCATCACATTATAAATGGATCAGTTGTTGACGGTTGTAGACTTTCAGTAACATAATTCTCATTTTATTCGTAGGCTTACAGAAAAATTCTCCCTATCGAGATATTTTTACATTAGCGGCATTAAGACTACAGGAATCTGGTGAGACAGAAAGATTACGACAAAAATGGTGGACTCAGCGTTCTCAATGTTCAGTTGTTGCTACATCTAGTGCTGATGCAGTTGAATTACAGGTGGGCAACGTTGCAGGTGTATTTATGGTCCTTGCTGGTGGCTTGATGTTGGCAACTCTATTTCTTATTATCGAACTGGTTATGTTTAAATTTTGTGAATCTCGCGCAAATAACCAGCGGACAATCAGACGTGTTCGTACTTCAATCATAATGCTATCAATTCATTATTCATTTCTATCTTATCTAAAATAAGTTATAATTGATAGTAATGACAATTAATGACTATTCCTATTTTTGATTTTAGAGAAATGAAGAATTAAATAGATCTGAATCAGGTCAAAATAATTTGGAATTTACTACTAGAAGCGAATCATCCAAATTAAGCCAAAATAATATTGCCTATGCTACCAGAACCAATAAAGTTAAATCAGTAAATTCTACCGACCTTTCGGAAATGATAACTAAACCTTCATCGTTGGGCCGCGATAATCGATCTTATAGTGACAACGATGTCAAAAAGCAGCAAGAGAATGATCGTAAATCAGAAACTGACCAACAACCAACTTCGACAGATCGACCCAAATCACAATTAACATCGACGAGTACGACTACTTCTGACTCACATAACCTCCCTGTTAGTTCTATCAGTAATAGTAAAATAATGTTTGTACATAATGATGATGATTCGATTTAA

Plasmid containing codon-optimized *Trichoplax adhaerans* AKDF^19393^ iGluR sequence

ATTTAGGTGACACTATAG, SP6 transcription promoter

AAGCTT, GGATCC, GAATTC, restriction sites

GCTTG etc, beta globin 5’ UTR and 3’ UTR

ACC, partial Kozak consensus sequence

TAA, stop codon

AAAAA etc, poly(A) sequence

CGTCAG, etc, remaining plasmid

CGTCAGCGGGTGTTGGCGGGTGTCGGGGCTGGCTTAACTATGCGGCATCAGAGCAGATTGTACTGAGAGTGCACCATTCGACGCTCTCCCTTATGCGACTCCTGCATTAGGAAGCAGCCCAGTAGTAGGTTGAGGCCGTTGAGCACCGCCGCCGCAAGGAATGGTGCATGCAAGGAGATGGCGCCCAACAGTCCCCCGGCCACGGGGCCTGCCACCATACCCACGCCGAAACAAGCGTTCATGAGCCCGAAGTGGCGAGCCCGATCTTCCCCATCGGTGATGTCGGCGATATAGGCGCCAGCAACCGCACCTGTGGCGCCGGTGATGCCGGCCACGATGCGTCCGGCGTAGAGGATCTGGCTAGCGATGACCCTGCTGATTGGTTCGCTGACCATTTCCGGGTGCGGGACGGCGTTACCAGAAACTCAGAAGGTTCGTCCAACCAAACCGATTCTGACGGCAGTTTACGAGAGAGATGATAGGGTCTGCTTCAGTAAGCCAGATGCTACACAATTAGGCTTGTACATATTGTCGTTAGAACGCGGCTACAATTAATACATAACCTTATGTATCATACACATACGATTTAGGTGACACTATAGAATACAAGCTTGCTTGTTCTTTTTGCAGAAGCTCAGAATAAACGCTCAACTTTGGCGTCGACACCATGCCTATATCCAAATTGGTTATCCTGCTTTACATTCTTATTTTCGTCTCCGCTGAGGAAATTCCAATCGTAAAAGTAAATCATAATGTCATTAAAATCGGCGTTTTTATGGAAAAATCCTCCATAGAGGGACGAACCGTCAGAGAAACCATTGATTTCATCAACAAGAATAAAGTCTATCCTTTACAGAATGCTACCCTGACGCCTGTCTTCCAGGCTATCGATAAAGACTCCCCTTACGTCGTTTCTCGTCAAGTGTGCCAATTTATAGAAAATGGTATTGTAGCTATTATTGGGTCTCTGCGAAATCCTCAGTCAATGGCAATTGCTGAATCTATTTTCTATCATTATACAATTCCTCATCTGGCTGTCAATTCCGTATTGCACTCTAAATATATAGGTAGCGAATCTTATAGTTTCCACCTGAAGCCATCCAAGCTGGCGGAGTTGGAAAAAATCTTCACCATCTTAGAAATTCACGAATATCGTAGAATTGCAGTTCTATACGATCAGACGTGTGATAGCGGAGAGTTGGTTAGTGTGTTACAGCTTGCACATCGCCGAGAAGTCAATGTCCAACTGGAAGAAATGTCTGATTGGAATTATACCACCAAGTCCGTTGCTATCTCAATTATAATTAATCGCATTAATCGCGCTGATATCAGAGTGGCTTATCTGTGTGTCGATCGTACCCTTATTGATGATATTGTTACATACGTCGTCCGTCAAGACATGCATACTTTTAAACATACTTGGATTATTAAAGACGAGGTTTTTCAGGATGCTGAGGATGATTTTGCTTCACTGAAGCGTTTATACAGTATAGGTTACTCCCCCGATACGAAGATGGAACAAAAGTGGAAGAATGTCAGAGTCGAACAAGCTATCTCGTATGATGCTGTCCATATTATTGCAAAAGCTATCAGTTCCACTATCCAATTAAATCAATTTCGTCACCAGAAGATTAATTGCAATCGTCATCCTCTGATCCCACTTACATCTGGACATGCTATCATGCGTCAGGTCAAAAAGCTTTATTTCTCTGGAGCTACGGGTAAGATTGAGTTCAATGATCGCGGAGAACGTCTGCATTGGAATCCTAAGATACATGATGTTCATAGTCGTATGAAGCAATCTTCGATCTCTAAGCGTGAGAAGCGTAGTGTACTTGCAGCTTCTGAAAGTAATGATAATGGTGAATCTTCGAATCCTTTAAGTCCCCAGAATTTACAGTTTATGGCCAATGAAACTCTTCGAATTACCACCGTGATAGATGCACCGTTTATCATGTACAAGCCCGAAAATCAACGTGGGCCTGGAATCGATGCTTTCGAGGGCTATTGCATTGACCTTCTGAAGTTACTGGTTGCTGAACTAGGATTCAACTTCACCATTAGTTTAGAACCTAGTAATATTTTTGGAGAACTTTTAGAAGACGGCAATTGGGACGGTATGGTAGGCCAGCTGGCTCGTAGGGAAATGGACATAGCTATCGCTCCATTATCTATTACGTCGTTTAGACAGTCCGTCATTGATTTTACCAAACCGTTTATGCAGTCAGGATTGCAGCTGTTAATTTCCAAGCCCAATCGTAATGCGGACTTTTTCGCTTTTCTGAGACCATTCTCACCGCTTGTTTGGATACTTATCGCTCTGGCTTTAATTTTTGTCGGAGTGGTGATGTATTTATTAGATCAATATTCTCCGGTCGATTTGGATTATGGAAATTGCTTCCCTGGATTGCGTTCTCGTTTTGGAAGTTCTAAAAAGAAAACTCGACTTAAGAAAGTTCTGCCTCGTAATGCTCTGCCTCGTGTCAAGTTTAACTTAGCGAACAGTTTCTGGTTTGCTATTGGTTCTTACTTACAACAAGGAGTTGAGGCTTATCCTCGATCTATCTCCGGACGTTTATTAGCTACATTTTATTGGTTCTTCGTTATGGTTATGGTCGCAAGCTATACAGCGAATATGGCTGCTTTTCTTACCACCTCTAATCTTGCTGTCCCGGTATCTTCCGTCCAAGACCTGACCAAGCAAAGCGCTATCGATTATGGAACCGTTATCGATTCCGAGGCTATGGATTTTTTAAAGTCATCGACCGAAACAGTCTACAGAAACGCTTGGTCGCATATCCTGGAAAATTCAAAGACTTCCCTGCTCAATTCATCCCTCGTCGGTATAAACAAAGTCCGTCGATCCAAGGGGAAGTACGCTTTCATTTGGTATTACCCTGAACTAGAATATGCAGCTCTGCGAAAACCTTGCGATGTTATGGTAGTCGGTTCCAAATTCGATCTACGCGGCTTTGGAATTGGTTTACAGAAGAACTCTCCATATAGAGATGTCTTTACCTTAGCTGCCTTAAAATTGCAAGAAAGAGGAGAAACAGAAAAGCTGCGTGCTAAGTGGTGGGATCAGCAGACTCAGTGTCCTCCTTCAACGTCTAGCGCAGCTGACGCAGTTGAATTGGGGGTCAACAATGTCGCTGGTATTTTCATGGTGCTTGGTGGTGCATTGATGTTGGCTACCATCCTTCTGATCATTGAACTTTTCATGTTCAAATTTTGCGTCTCACGTGCCAGTTCCAAGCGAACCATGAGACAAAGAAATGGTAACTTGATTAATGCTGAAATGGGTGAAAATAAGTTAGAATTTACTTCTCGTAAAGTCAGCTCTGGTAATATCAATGCTCTGGATATACCCAGAGCGACCACCACAGTTCCTGCAAGTACCGTTCATGATAACCAGGGTTACATATCCAACGAAATACAGACCGAAGACAATCAACATATACAAATCGATCAGCGATCCCCAACCCAAGCTGACTCTCCATCCACCCAGACAGACACCACCAATTCCGACACACAAAACCACGGTATTAGTCCTACTAATAATAACAATTGCACATTTAGGCAAAACTAAGGATCCGGTTACCACTAAACCAGCCTCAAGAACACCCGAATGGAGTCTCTAAGCTACATAATACCAACTTACACTTTACAAAATGTTGTCCCCCAAAATGTAGCCATTCGTATCTGCTCCTAATAAAAAGAAAGTTTCTTCACATTCTAAAAAAAAAAAAAAAAAAAAAAAAAAAAAACGAATTCGTAATCATGTCATAGCTGTTTCCTGTGTGAAATTGTTATCCGCTCACAATTCCACACAACATACGAGCCGGAAGCATAAAGTGTAAAGCCTGGGGTGCCTAATGAGTGAGCTAACTCACATTAATTGCGTTGCGCTCACTGCCCGCTTTCCAGTCGGGAAACCTGTCGTGCCAGCTGCATTAATGAATCGGCCAACGCGCGGGGAGAGGCGGTTTGCGTATTGGGCGCTCTTCCGCTTCCTCGCTCACTGACTCGCTGCGCTCGGTCGTTCGGCTGCGGCGAGCGGTATCAGCTCACTCAAAGGCGGTAATACGGTTATCCACAGAATCAGGGGATAACGCAGGAAAGAACATGTGAGCAAAAGGCCAGCAAAAGGCCAGGAACCGTAAAAAGGCCGCGTTGCTGGCGTTTTTCCATAGGCTCCGCCCCCCTGACGAGCATCACAAAAATCGACGCTCAAGTCAGAGGTGGCGAAACCCGACAGGACTATAAAGATACCAGGCGTTTCCCCCTGGAAGCTCCCTCGTGCGCTCTCCTGTTCCGACCCTGCCGCTTACCGGATACCTGTCCGCCTTTCTCCCTTCGGGAAGCGTGGCGCTTTCTCATAGCTCACGCTGTAGGTATCTCAGTTCGGTGTAGGTCGTTCGCTCCAAGCTGGGCTGTGTGCACGAACCCCCCGTTCAGCCCGACCGCTGCGCCTTATCCGGTAACTATCGTCTTGAGTCCAACCCGGTAAGACACGACTTATCGCCACTGGCAGCAGCCACTGGTAACAGGATTAGCAGAGCGAGGTATGTAGGCGGTGCTACAGAGTTCTTGAAGTGGTGGCCTAACTACGGCTACACTAGAAGAACAGTATTTGGTATCTGCGCTCTGCTGAAGCCAGTTACCTTCGGAAAAAGAGTTGGTAGCTCTTGATCCGGCAAACAAACCACCGCTGGTAGCGGTGGTTTTTTTGTTTGCAAGCAGCAGATTACGCGCAGAAAAAAAGGATCTCAAGAAGATCCTTTGATCTTTTCTACGGGGTCTGACGCTCAGTGGAACGAAAACTCACGTTAAGGGATTTTGGTCATGAGATTATCAAAAAGGATCTTCACCTAGATCCTTTTAAATTAAAAATGAAGTTTTAAATCAATCTAAAGTATATATGAGTAAACTTGGTCTGACAGTTACCAATGCTTAATCAGTGAGGCACCTATCTCAGCGATCTGTCTATTTCGTTCATCCATAGTTGCCTGACTCCCCGTCGTGTAGATAACTACGATACGGGAGGGCTTACCATCTGGCCCCAGTGCTGCAATGATACCGCGAGACCCACGCTCACCGGCTCCAGATTTATCAGCAATAAACCAGCCAGCCGGAAGGGCCGAGCGCAGAAGTGGTCCTGCAACTTTATCCGCCTCCATCCAGTCTATTAATTGTTGCCGGGAAGCTAGAGTAAGTAGTTCGCCAGTTAATAGTTTGCGCAACGTTGTTGCCATTGCTACAGGCATCGTGGTGTCACGCTCGTCGTTTGGTATGGCTTCATTCAGCTCCGGTTCCCAACGATCAAGGCGAGTTACATGATCCCCCATGTTGTGCAAAAAAGCGGTTAGCTCCTTCGGTCCTCCGATCGTTGTCAGAAGTAAGTTGGCCGCAGTGTTATCACTCATGGTTATGGCAGCACTGCATAATTCTCTTACTGTCATGCCATCCGTAAGATGCTTTTCTGTGACTGGTGAGTACTCAACCAAGTCATTCTGAGAATAGTGTATGCGGCGACCGAGTTGCTCTTGCCCGGCGTCAATACGGGATAATACCGCGCCACATAGCAGAACTTTAAAAGTGCTCATCATTGGAAAACGTTCTTCGGGGCGAAAACTCTCAAGGATCTTACCGCTGTTGAGATCCAGTTCGATGTAACCCACTCGTGCACCCAACTGATCTTCAGCATCTTTTACTTTCACCAGCGTTTCTGGGTGAGCAAAAACAGGAAGGCAAAATGCCGCAAAAAAGGGAATAAGGGCGACACGGAAATGTTGAATACTCATACTCTTCCTTTTTCAATATTATTGAAGCATTTATCAGGGTTATTGTCTCATGAGCGGATACATATTTGAATGTATTTAGAAAAATAAACAAATAGGGGTTCCGCGCACATTTCCCCGAAAAGTGCCACCTGACGTCTAAGAAACCATTATTATCATGACATTAACCTATAAAAATAGGCGTATCACGAGGCCCTTTCGTCTCGCGCGTTTCGGTGATGACGGTGAAAACCTCTGACACATGCAGCTCCCGGAGACGGTCACAGCTTGTCTGTAAGCGGATGCCGGGAGCAGACAAGCCCGTCAGGGCG

**Supplemental information references**

1.	M. E. Saks *et al.*, An engineered Tetrahymena tRNAGln for in vivo incorporation of unnatural amino acids into proteins by nonsense suppression. *J Biol Chem* **271**, 23169-23175 (1996).

2.	M. Eitel *et al.*, Comparative genomics and the nature of placozoan species. *PLoS Biol* **16**, e2005359 (2018).

3.	A. I. Sobolevsky, M. P. Rosconi, E. Gouaux, X-ray structure, symmetry and mechanism of an AMPA-subtype glutamate receptor. *Nature* **462**, 745-756 (2009).
